# Permeability profiling of all 13 Arabidopsis PIP aquaporins using a high throughput yeast approach

**DOI:** 10.1101/2021.05.09.443061

**Authors:** Michael Groszmann, Annamaria De Rosa, Weihua Chen, Jiaen Qiu, Samantha A McGaughey, Caitlin S. Byrt, John R Evans

## Abstract

Plant aquaporins have many more functions than just transporting water. Within the diversity of plant aquaporins are isoforms capable of transporting signaling molecules, nutrients, metalloids and gases. It is established that aquaporin substrate discrimination depends on combinations of factors such as solute size, pore size and polarity, and post-translational protein modifications. But our understanding of the relationships between variation in aquaporin structures and the implications for permeability is limited. High-throughput yeast-based assays were developed to assess diverse substrate permeabilities to water, H_2_O_2_, boric acid, urea and Na^+^. All 13 plasma membrane intrinsic proteins (PIPs) from Arabidopsis (AtPIPs) were permeable to both water and H_2_O_2_, although their effectiveness varied, and none were permeable to urea. AtPIP2 isoforms were more permeable to water than AtPIP1s, while AtPIP1s were more efficient at transporting H_2_O_2_ with AtPIP1;3 and AtPIP1;4 being the most permeable. Among the AtPIP2s, AtPIP2;2 and AtPIP2;7 were also permeable to boric acid and Na^+^. Linking AtPIP substrate profiles with phylogenetics and gene expression data enabled us to align substrate preferences with known biological roles of AtPIPs and importantly guide towards unidentified roles hidden by functional redundancy at key developmental stages and within tissue types. This analysis positions us to more strategically test *in planta* physiological roles of AtPIPs in order to unravel their complex contributions to the transport of important substrates, and secondly, to resolve links between aquaporin protein structure, substrate discrimination, and transport efficiency.

**One sentence summary:** Yeast based high throughput assays were developed to assess the permeability of each Arabidopsis PIP aquaporin isoform to water, H_2_O_2_, boric acid, urea and sodium.

## Introduction

Aquaporins (AQPs) are membrane intrinsic proteins (MIPs) and constitute a major family of channel proteins found across all phylogenetic kingdoms (Chaumont and Tyerman, 2017). AQP monomers form a characteristic hour-glass membrane-spanning pore that differ in aperture and residue composition which determines their particular substrate selectivity and permeabilities. Four AQP monomers assemble to form tetrameric complexes which create a fifth central pore that has been implicated for the movement of CO_2_ (Kaldenhoff *et al*., 2014) and ions (Yu *et al*., 2006) across membranes.

The AQP gene family has diversified to the greatest extent in plants. This may reflect greater duplication rates of plant genomes and the adaptation potential AQPs provide for a sessile lifestyle. Genomes of Angiosperm species commonly harbour between 30-50 isoforms, with extremes of 84 and 121 in tobacco and canola, respectively (Groszmann *et al*., 2017, Sonah *et al*., 2017, De Rosa *et al*., 2020, Groszmann *et al*., 2021). Of the 13 AQP subfamilies recognised in the plant kingdom, five subfamilies predominate in the angiosperms (PIPs, TIPs, NIPs, SIPs, and XIPs)(Laloux *et al*., 2018). Each subfamily is generally characterised by sequence composition, a tendency to localise to different subcellular membranes, and transport different sets of substrates. Key pore features such as the dual Asn-Pro-Ala (NPA) motifs, the aromatic/Arginine (ar/R) filter and Froger’s position have been associated with broader substrate selectivity (e.g. water vs. urea). However, gaining a more nuanced understanding of signatures related to substrate selectivity, transport efficiency and substrate exclusivity between isoforms requires more detailed characterisation. While a single AQP isoform can permeate a variety of substrates, surprisingly few have been surveyed for multiple substrates in parallel under similar conditions to establish comparative transport profiles.

Plant AQPs are implicated in numerous physiological processes including: water relations, organ growth, fertilisation, seed development and germination, abiotic stress responses, defence signalling, nutrient uptake and tolerance, and photosynthesis (Chaumont and Tyerman, 2017). Plant AQPs are permeable to many substrates indispensable for plant growth such as, water, CO_2_ and nitrogen (NH_3_/NH_4_^+^, urea and nitrate); micronutrients (boric acid and silicic acid) and other metalloids; signalling molecules hydrogen peroxide (H_2_O_2_) and nitric oxide (NO); O_2_ and lactic acid to cope with anoxic stress; and key nutrients such as potassium (Chaumont and Tyerman, 2017, Qiu *et al*., 2020, Singh *et al*., 2020). The diverse substrate specificities and involvement in key plant processes make AQPs interesting targets for engineering more resilient and productive crops (Afzal *et al*., 2016, Singh *et al*., 2020), and for use in industrial filtration applications (Tang *et al*., 2015, Hélix-Nielsen, 2018, Jafarinejad, 2020).

The increasing number of curated *AQP* gene families offers a rich source of isoform variation information. Having a high-throughput permeability assessment system for testing different isoforms would enable the building of a catalogue of information about their substrate profiles. The substrate selectivity and functional capacity of AQPs are routinely assessed in heterologous systems such as oocytes, liposomes, artificial membranes, and yeast (Madeira *et al*., 2016). Most of these systems and assays require specialized equipment (e.g. stopped-flow spectrophotometer), or complicated setups (e.g. artificial polymer membranes), or are labor intensive (e.g. *Xenopus laevis* oocytes), which preclude their use for high-throughput applications. By contrast, yeast offer a simple and versatile host for the heterologous production of aquaporins (Öberg *et al*., 2009, Bill, 2014), with which to test different substrates.

The diversity of well characterized mutant strains of *S. cerevisiae* enables bespoke optimization for screening specific substrate permeabilities of heterologously expressed AQPs. Mutant strains are available that are sensitive to a given cytotoxic agent, or where native transporters for compounds essential for growth that are not functional have been replaced by alternative uptake routes associated with the heterologously expressed AQP.

Altered sensitivity of AQP-expressing yeast can be detected through cell dilution spot tests for colony formation on solid medium containing the test substrate. While this traditional method is more accessible, it has several drawbacks including being poorly quantitative (Hung *et al*., 2018). Real-time optical density (OD) monitoring of yeast micro-volume cultures (< 300μl) can overcome the limitations of agar-based spot assays. They are particularly suitable for detecting small phenotypic changes in yeast population growth and are a well-established method for monitoring responses to chemical treatments (Warringer and Blomberg, 2003, Toussaint *et al*., 2006, Marešová and Sychrová, 2007).

Here, we establish a qualitative and quantitative methodological framework, involving a high-throughput micro-cultivation-based yeast system, to functionally characterize AQP transport selectivity and capacity. We applied these methods to all 13 members of the Arabidopsis PIP aquaporin family (AtPIPs), determining their permeabilities to water, hydrogen peroxide, boric acid, urea and sodium. This type of approach could be used to efficiently catalogue the transport capacity of a large number of AQPs to clarify their biological roles in plants. It also has the potential to help decipher the nuanced characteristics of transport selectivity and efficiency necessary for future engineering of AQPs for specific biotechnological applications.

## Results

### Developing high-throughput micro-volume yeast culturing assays to assess aquaporin function

#### Optimizing conditions for reproducible growth curves

We established a high-throughput *Saccharomyces cerevisiae* (yeast) micro-cultivation (200 μl) method using 96-well plates. The micro-cultures were incubated in a plate reader with versatile control over temperature, shaking, and OD reading modes. We optimized these parameters to find conditions that generated repeatable growth curves (Fig. 1A; see Supplemental Materials and Methods for details). We observed that micro-volume cultures tended to aggregate and sediment in wells regardless of the shaking intensity. Sedimentation was managed using a double orbital shaking mode which dispersed yeast evenly across the bottom of the well and recording OD as an average of multiple measurements at distinct points around each well using the well scanning mode on the plate reader.

**Figure 1.**
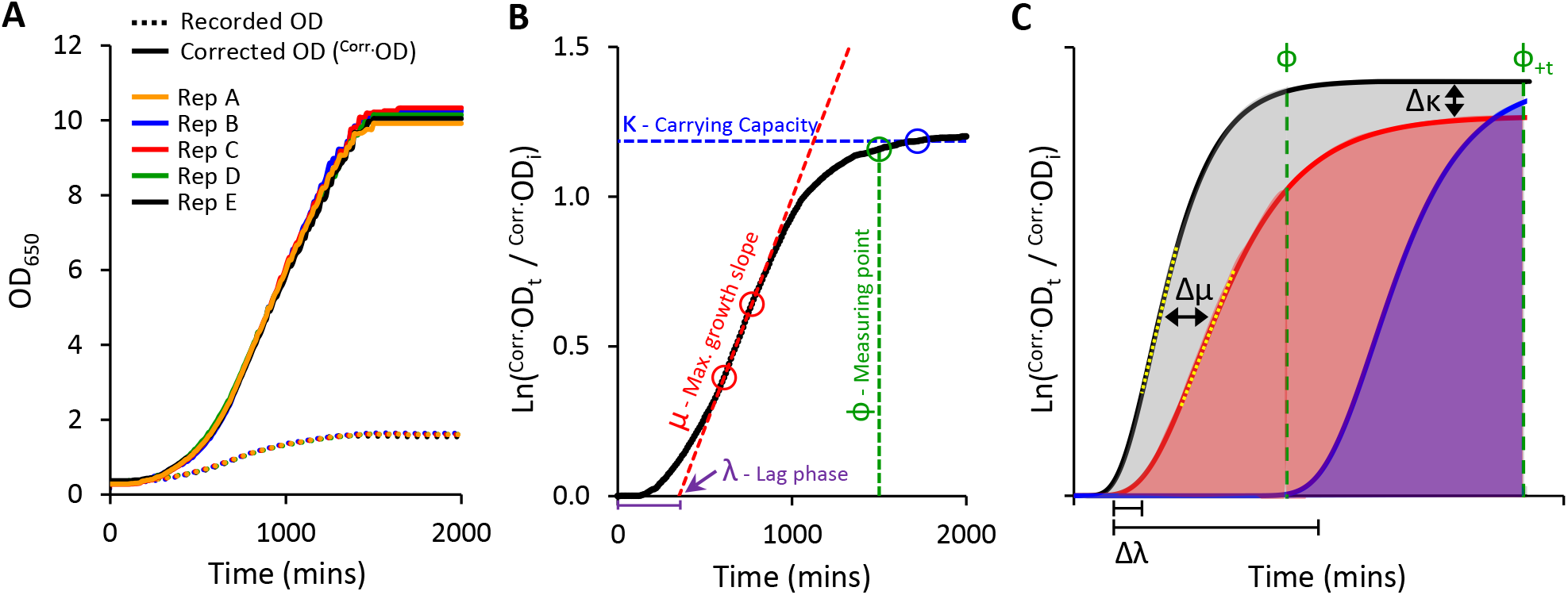
Yeast micro-cultivation setup and growth data outputs. **A**, Optimised micro-cultivation conditions produce repeatable growth curves of replicate cultures spaced across a 96-well plate. The growth curves of recorded OD values are compressed due to the progressive non-linear response of optical detection. Applying a calibration function produces corrected OD values (^Corr.^OD) and a more accurate representative growth curve. **B**, A yeast population growth curve (Ln ^Corr.^Od_t_ / ^Corr.^Od_i_) depicting the three major derived growth traits (λ, μ, and κ) and the dynamic standardizing measuring point, Phi (ϕ). **C**, Conceptual examples demonstrating the use of Area Under the Curve (AUC) as a measure of cumulative growth differences. Untreated yeast population growth (black) and two treatment growth scenarios (blue and red). Φ is allocated to the untreated growth curve. The red curve shows a slightly longer lag phase (Δλ), reduced maximum rate of growth (Δμ; differences between yellow dotted tangent lines), and lower carrying capacity (Δκ), captured as a substantially reduced AUC (shading) than that of the untreated black curve. The blue curve shows a longer lag phase, but growth rate and carrying capacity similar to untreated. No AUC is detected at ϕ, but AUC can be detected by shifting to ϕ_+t_ (note: ΔAUC will be less (underestimated) when using ϕ_+t_ as control population has ceased growing).

#### Adjusting for non-linearity of OD measurements at high cell density

Growing yeast cultures quickly achieve densities that far exceed saturation limits of optical detection in spectrophotometers (Fig. 1A) (Stevenson *et al*., 2016). This severely underestimates ‘true’ ODs at higher cell densities, resulting in compressed growth curves and systematic distortion of extracted fitness components required to evaluate culture health and growth (Warringer and Blomberg, 2003, Fernandez-Ricaud *et al*., 2016).

We compared ‘recorded’ ODs against ‘true’ ODs calculated from dilution factors. A single polynomial function described the relationship between ‘recorded’ and ‘true’ OD datasets that was valid for all of the strains of *S. cerevisiae* used in this study (R^2^ > 0.99; Supplemental Fig. S1). Applying this calibration function to calculate corrected OD values (^Corr.^OD), improved the resolution of key derived growth characteristics: initial lag phase (λ), maximum growth rate (μ), and final carrying capacity or biomass yield (κ) (Fig. 1B).

#### Establishing the Phi (ϕ) measuring point and AUC value

To simplify the phenotyping, we calculated Area Under the Curve (AUC) as a single all-encompassing parameter that captured λ, μ and κ (Fig. 1C). We observed that heterologous expression of AtPIPs can differentially alter yeast growth traits independent of chemical treatment (Supplemental Table S1). This may occur to an even greater extent when assessing more diverse AQP isoforms. Altered inherent growth would mean yeast cultures mature at different rates, thereby complicating the evaluation of growth differences, especially when measuring all cultures at a single time point. Measuring a given culture sub-set too soon potentially misses growth phenotypes arising from subtle responses to treatments. Measuring too late, and the rapidly growing control cultures have plateaued, allowing the slower growing treated cultures time to catch up and reduce the difference. To account for variation in culture maturity times, we implemented a dynamic standardizing measuring point termed Phi (ϕ), defined just prior to the stationary phase of log transformed growth curves, at the point the population growth rate drops below 5% of maximum (Fig. 1B). ϕ is established on the best growing culture for a given AQP set (Fig. 1C), i.e. the untreated control when evaluating cytotoxic compounds (e.g. H_2_O_2_), or the culture with the highest supplementation of essential nutrient when examining growth requiring agents (e.g. urea). AUCs for all cultures were calculated from the start of cultivation until ϕ (Fig. 1C), with AUC_treated_/AUC_control_ providing relative differences in growth (ΔAUC). In our routine conditions, all control cultures reached and remained in stationary phase for an extended period of time. As such, ϕ can be shifted (ϕ_+t_) in order to capture additional data from treated cultures that grow very slowly; with an understanding that ΔAUC will be underestimated because the control culture plateaued earlier (Fig. 1C). Once ΔAUC values are established for each AQP, they are compared between AQPs to rank transport efficiencies.

### Heterologous AtPIP production in yeast

Having an abundance of AQP protein is the first essential requirement for robust functional evaluations and improves the detection limit in response to treatments. For example, the water permeability for AtPIP2;3 was assessed using two promoters, with greater freeze-thaw tolerance (a proxy for water permeability) achieved using the strong GPD promoter relative to the less active TPI1 promoter (Supplemental Fig. S2). To maximize the likelihood of high AtPIP production we (i) used high copy number plasmids with minimal load burdens on yeast growth, (ii) used a strong constitutive GPD promoter with complementing terminator, (iii) ensured codon usage compatibility between AtPIPs and yeast, and (iv) modified the Kozak sequence to enhance translational initiation (see Supplemental Materials and Methods). A parallel collection of *AtPIP-GFP* transgenes that differed only in the C-terminal GFP fusion compared to the expression vectors used in the functional assays, were used for evaluating heterologous AtPIP production *in vivo* and subcellular localization. All 13 AtPIP-GFP yeast lines repeatedly emitted strong GFP signal, indicating high AtPIP protein production (Supplemental Fig. S3), except for AtPIP1;4 which had 27% of the average fluorescence intensity. However, as described below, substrate transport associated with AtPIP1;4 was comparable with other AtPIP1 isoforms.

### Subcellular localization of AtPIPs in yeast

In addition to ample heterologous protein production, sufficient AtPIP needs to localize to the yeast plasma membrane (PM) in order to evaluate AQP-driven changes in substrate permeation into the cell. Sub-cellular localization of the AtPIPs was evaluated using confocal microscopy of AtPIP-GFP lines and compared against cytosolic (GFP only) and endoplasmic reticulum (ER; SEC63-RFP) markers (Fig. 2). Free GFP is cytosolically localized (Fig. 2A). The SEC63-RFP marker reveals the web-like ER network, with the prominent nuclear envelope ER domain (nER) and peripheral or cortical ER domain (cER) (Fig. 2B). The cER lies immediately adjacent to the plasma membrane but is discontinuous around the perimeter with discernible gaps distinguishing it from PM localisation (Fig. 2B). A sharp ring around the cell perimeter was seen for all 8 AtPIP2-GFP proteins, consistent with strong PM integration (Fig. 2, E, F, I, J, M, N, Q and R). By contrast, when expressed alone, the five AtPIP1-GFP proteins show dual localization consisting of a patchy peripheral ring and internal webs like the SEC63-RFP ER marker (Fig. 2, C, G, K, O and S), along with a distinct continuous ring around the periphery indicating PM localization, but less efficient than AtPIP2s.

**Figure 2.**
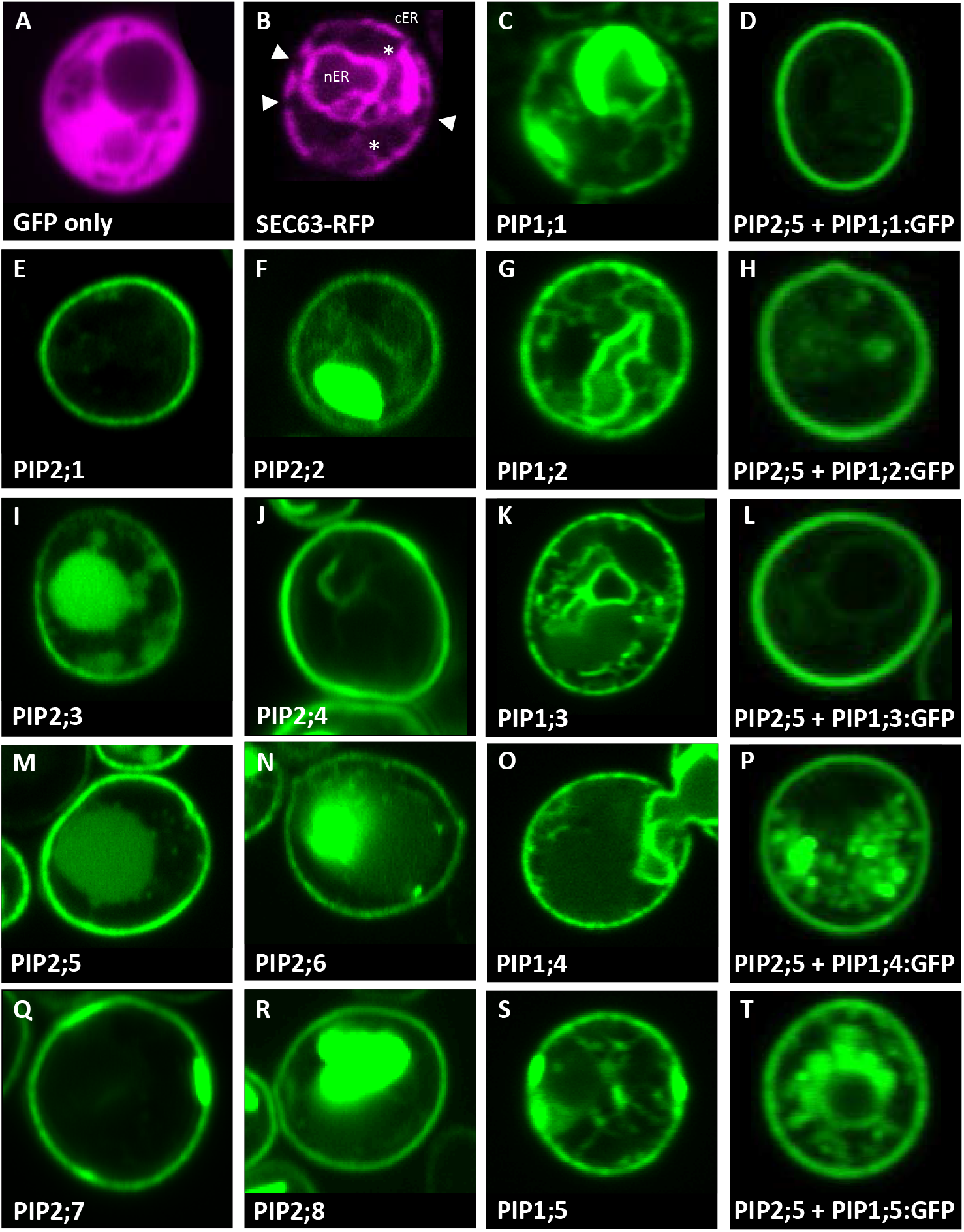
Sub-cellular localisation of AtPIPs in yeast. Confocal microscopy images of: **A**, an eGFP only control showing diffuse cytosolic localised signal. **B**, SEC63::RFP endoplasmic reticulum (ER) marker showing the prominent nuclear envelope ER domain (nER) and a peripheral or cortical ER domain (cER). The cER lies just beneath the plasma membrane but is not continuous around the perimeter with gaps distinguishing it from plasma membrane localisation (solid triangles). Cytoplasmic tubules link the two ER domains (*). **E, F, I, J, M, N, Q and R**, AtPIP2-eGFP proteins expressed alone predominantly localise in a distinct continuous ring of signal around the cell perimeter coinciding with the plasma membrane. In several cases, eGFP signal can also be detected in internal storage vacuoles. **C, G, K, O and S**, AtPIP1-eGFP proteins expressed alone localise to the nER, ER tubules and a patchy cER signal overlaying PM localisation. **D, H, L, P and T**, AtPIP1-eGFP proteins co-expressed with AtPIP2;5 with the majority of the fluorescence signal localised to the PM, similar to AtPIP2 proteins. Fluorescence signal false colored red for marker lines in A and B, and green for AtPIP-GFP lines in C-T.

### Co-expression with AtPIP2;5 enables AtPIP1s to more efficiently localize to the yeast PM

PIP2 proteins can interact and guide PIP1 proteins more efficiently to the PM (Jozefkowicz *et al*., 2017). The Yeast-two-Hybrid mating-based Split-Ubiquitin System (Y2H mbSUS; Fig. 3A) was used to screen an AtPIP interactome library. Yeast co-expressing the bait AtPIP2;5-CubPLV and any of the AtPIP1;1-Nub to AtPIP1;5-Nub prey proteins, activated the *lacZ* reporter ≥ 4-fold above background levels (Fig. 3B), demonstrating that AtPIP2;5 strongly interacted with each AtPIP1. Co-expression of *AtPIP2*;*5* with *GFP* tagged versions of *AtPIP1*;*1* to *1*;*5*, resulted in most of the fluorescence signal now being associated with the PM (Fig. 2, D, H, L, P and T). *AtPIP2*;*5* was chosen because, among the *AtPIP2s*, it showed moderate levels of apparent permeability to the tested substrates, enabling further improvements in permeability due to the co-expressed AtPIP1 isoforms to be observed.

**Figure 3.**
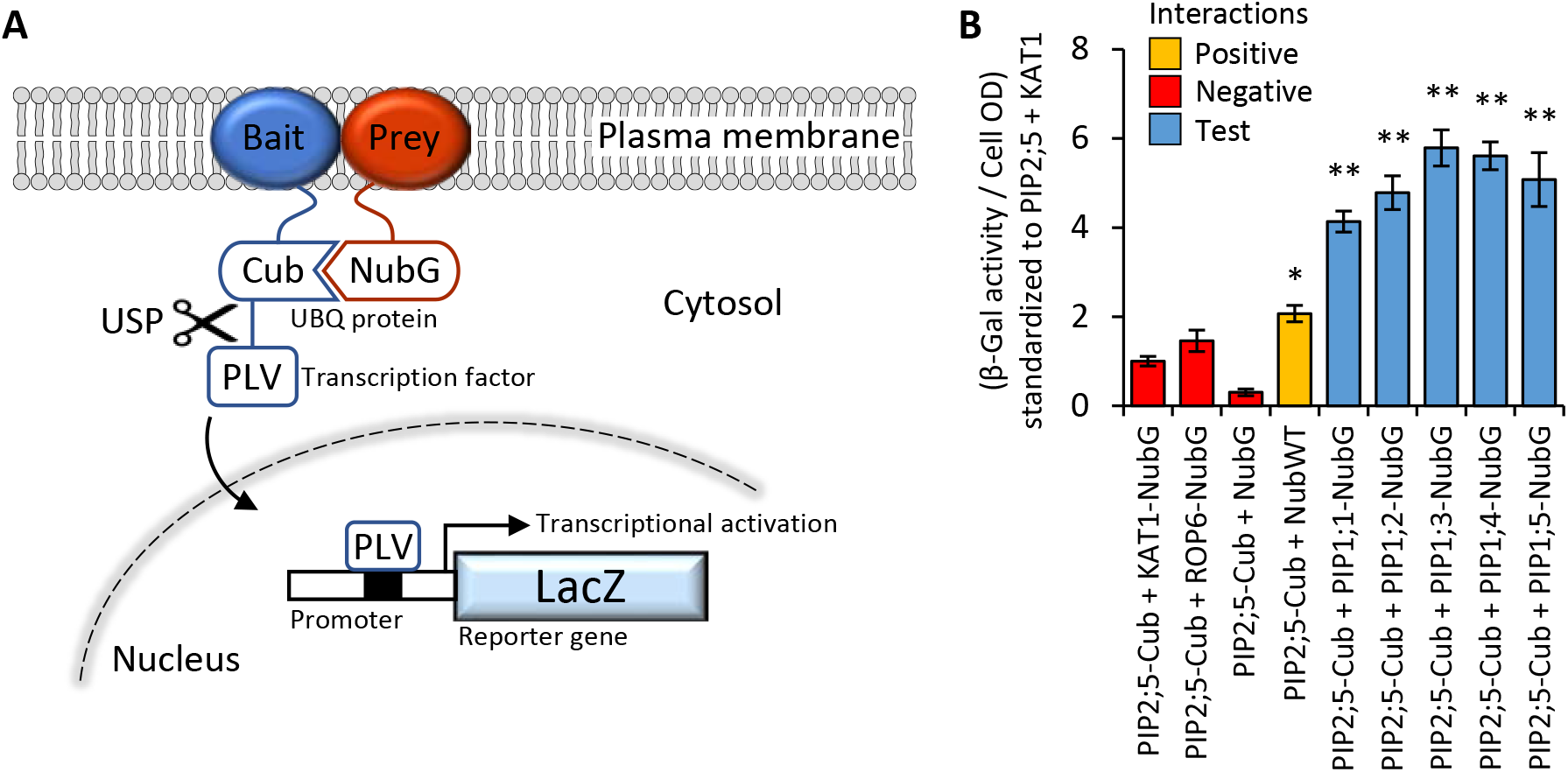
**A**, Illustration of mbSUS yeast two-hybrid system. The mutant N-terminal ubiquitin domain (NubG) and C-terminal ubiquitin domain (Cub) can reconstitute the full-length ubiquitin protein (UBQ) only when brought into close proximity via a membrane bound and interacting Bait and Prey protein combination. The reconstituted UBQ is recognised by Ubiquitin-Specific Proteases (USP), releasing the artificial transcription factor PLV (proteinA-LexA-VP16) that is translationally fused to the Cub domain. The freed PLV then enters the nucleus and activates the *LacZ* reporter gene that encodes for a β-galactosidase. **B**, AtPIP2;5 is capable of strong protein-protein interactions with each of the AtPIP1 isoforms. The intensity of the AtPIP2;5 (bait) and AtPIP1 (prey) interaction was assayed by measuring β-galactosidase activity via colorimetric monitoring of o-nitrophenyl-β-D-galactoside (ONPG) conversion to the yellow o-nitrophenol. Control lines: NubG (pNX35-DEST), a mutant Nub variant with low affinity for Cub. When linked to plasma membrane localizing Arabidopsis ROP6 or KAT1 proteins, it acts as a prey control reporting incidental UBQ reconstitution through simple random close insertion of abundantly produced membrane bound proteins. NubG expressed alone should not interact with Cub and negligible reporter activity was observed. NubWT (pNubWTXgate) is a soluble cytoplasmic localized N-terminal ubiquitin domain with a high affinity for Cub and acts as a positive control able to interact with the Cub domain of AtPIP2;5-Cub independent of bait interaction. The detected activity (orange) demonstrates that the Cub domain fused to AtPIP2;5 was accessible to Nub and USPs. Each of the AtPIP2;5 + AtPIP1 interactions (blue) significantly exceeded spurious background levels (red). All error bars are SEM. ANOVA post-hoc Fisher’s LSD versus ATPIP2;5 + KAT1, * p < 0.05, ** p < 0.01. N = 4 biological reps over 2 experimental runs.

### Characterizing AtPIP water permeability

The permeability of AtPIPs to water was tested using a rapid freeze-thaw assay adapted to our micro-cultivation setup. For wild type yeast carrying an empty vector, successive freeze-thaw treatments incrementally decreased ΔAUC (Supplemental Fig. S4, A and B). Freeze-thawing prolonged the lag phase (Supplemental Fig. S4C), consistent with a reduction in the viable cell count of the starting population, which delayed the detection of population growth. The sensitivity of the freeze-thaw assay was improved by using the *aquaporin* null mutant background (*aqy1 aqy2*), which is compromised in tolerance to rapid freeze-thaw events (Tanghe *et al*., 2002, Tanghe *et al*., 2004). Two freeze-thaw cycles were sufficient to essentially render the entire *aqy1 aqy2* starting population unviable (Supplemental Fig. S4, A-C). Heterologous expression of a water permeable AQP (*AtPIP2*;*1*) (Verdoucq *et al*., 2008), dramatically improved the tolerance of the *aqy1 aqy2* mutant to repeated freeze-thaw treatments (Supplemental Fig. S4, A-C).

Application of two freeze-thaw treatments to *aqy1 aqy2* yeast carrying one of the 13 AtPIP genes or an empty vector differentially affected the growth curves (Fig. 4A). All of the AtPIP2 proteins had sufficient capacity to transport water across the PM to confer freeze-thaw tolerance, but their effectiveness varied with AtPIP2;7 the most effective and AtPIP2;2 the least effective (Fig. 4B). At ϕ, growth was not detected for any *AtPIP1* expressing lines. Freeze-thaw tolerance associated with AtPIP1s was revealed by calculating AUC at ϕ + 1000 mins, but resolution between AtPIP2 isoforms was lost (Fig. 4C). The implied water transport capacity of AtPIP1s were substantially lower than the AtPIP2s.

**Figure 4.**
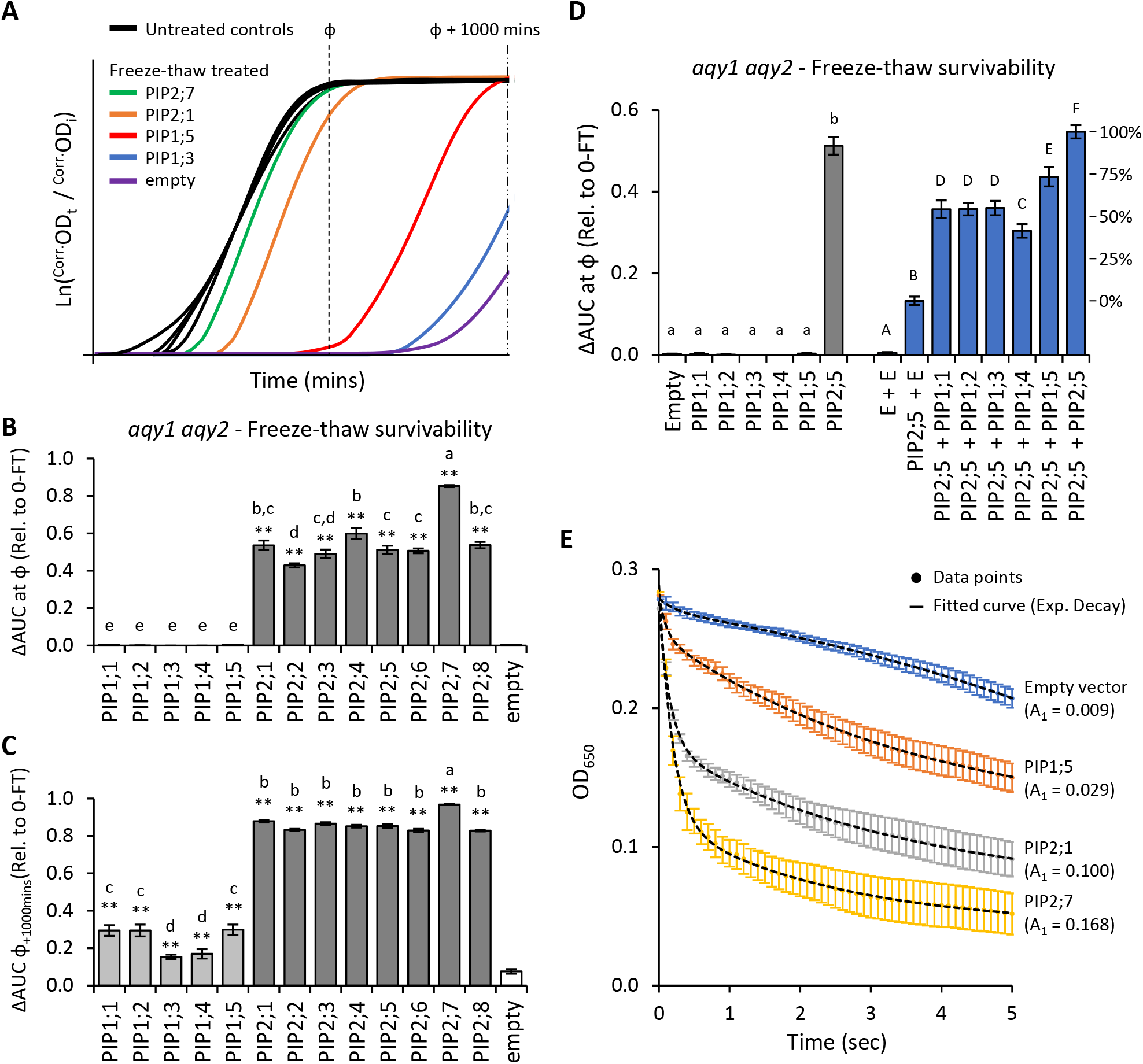
Water permeability assays using two freeze-thaw cycles with yeast expressing different AQP genes. **A**, Illustrative growth curves for untreated controls and following two freeze-thaw cycles. **B**, Relative AUC for the 13 AtPIP isoforms, calculated with ϕ. **C**, Relative AUC after extended growth with AUC calculated at ϕ+1000. **D**, Relative AUC for AtPIP1s expressed singly or co-expressed with AtPIP2;5. **E**, Change in OD of yeast spheroplast suspensions following osmotic shock. The contribution of the rapid initial phase (value in parentheses) reflects the permeability derived from fitted two-phase exponential curves; empty vector: y = [0.00881 x *e*^(-x/0.243)^] + [-0.05398 x *e*^(-x/-6.47128)^]; AtPIP1;5: y = [0.02937 x *e*^(-x/0.09966)^] + [0.13874 x *e*^(-x/3.76055)^]; AtPIP2;1: y = [0.10037 x *e*^(-x/0.15797)^] + [0.10763 x *e*^(-x/3.51469)^]; AtPIP2;7: y = [0.16814 x *e*^(-x/0.18973)^] + [0.07791 x *e*^(-x/2.43538)^]. All error bars are SEM. For B-C, asterisks indicate statistical difference from empty vector control, ANOVA with Fishers LSD test (* *P* < 0.05; ** *P* < 0.01); letters denote different statistical rankings, ANOVA with Tukey’s test (*P* < 0.05). For D, letters denote different statistical groupings, lowercase among single expressed and uppercase among co-expressed AtPIP yeast lines, ANOVA with Tukey’s test (*P* < 0.05). N = 12 (AtPIP1s) and 8 (AtPIP2s) across 4 experimental runs for B and C. N = 6 across 3 experimental runs for D. N = 6 across 2 experimental runs for E.

Water permeability of AtPIP1s was further assessed by increasing their abundance in the PM through co-expression with *AtPIP2*;*5*. Yeast co-transformed with *AtPIP2*;*5* + *Empty* vector served as a base-level control, with less freeze-thaw tolerance than yeast carrying the *AtPIP2*;*5* vector alone or co-expressing two copies of *AtPIP2*;*5* (Figure 4D). This is consistent with *AtPIP2*;*5* + *Empty* vector yeast having reduced expression of *AtPIP2*;*5* as only half the plasmid load carries *AtPIP2*;*5*. Co-expression of *AtPIP1*;*1*, *1*;*2*, *1*;*3*, *1*;*4* or *1*;*5* with *AtPIP2*;*5* substantially improved freeze-thaw survivorship over the *AtPIP2*;*5* + *Empty* vector control, being from ~40-75% as effective as AtPIP2;5 (i.e. AtPIP2;5 + AtPIP2;5; Fig. 4D). This revealed that AtPIP1 isoforms have significant capacity to transport water, but are less efficient than AtPIP2s.

Water permeability was also assessed using the traditional, but more laborious, yeast spheroplast bursting method (Fig. 4E). Water transport efficiencies were ranked AtPIP2;7 > AtPIP2;1 > AtPIP1;5 > empty, matching the order and approximate magnitude of differences obtained from the freeze-thaw assay. The consistency in results from the two methods validated assessment of water permeability across the AtPIP family using the freeze-thaw assay which provided a qualitative and quantitative platform to rapidly evaluate water transport capacity of AQPs.

### Characterization of AtPIP H_2_O_2_ permeability

Hydrogen peroxide (H_2_O_2_) treatments impaired growth of the empty vector *aqy1 aqy2* yeast (Fig. 5A), impacting all three growth traits (λ, μ, and κ; Supplemental Fig. S5). The effects were more prominent when using the *skn7* yeast which is compromised in its antioxidant buffering capacity (Fig. 5A; Supplemental Fig. S5). 0.5mM and 1mM H_2_O_2_ were chosen as treatment concentrations as they occur at the commencement of pronounced growth inhibition on the dose response curves (Figure 5B).

**Figure 5.**
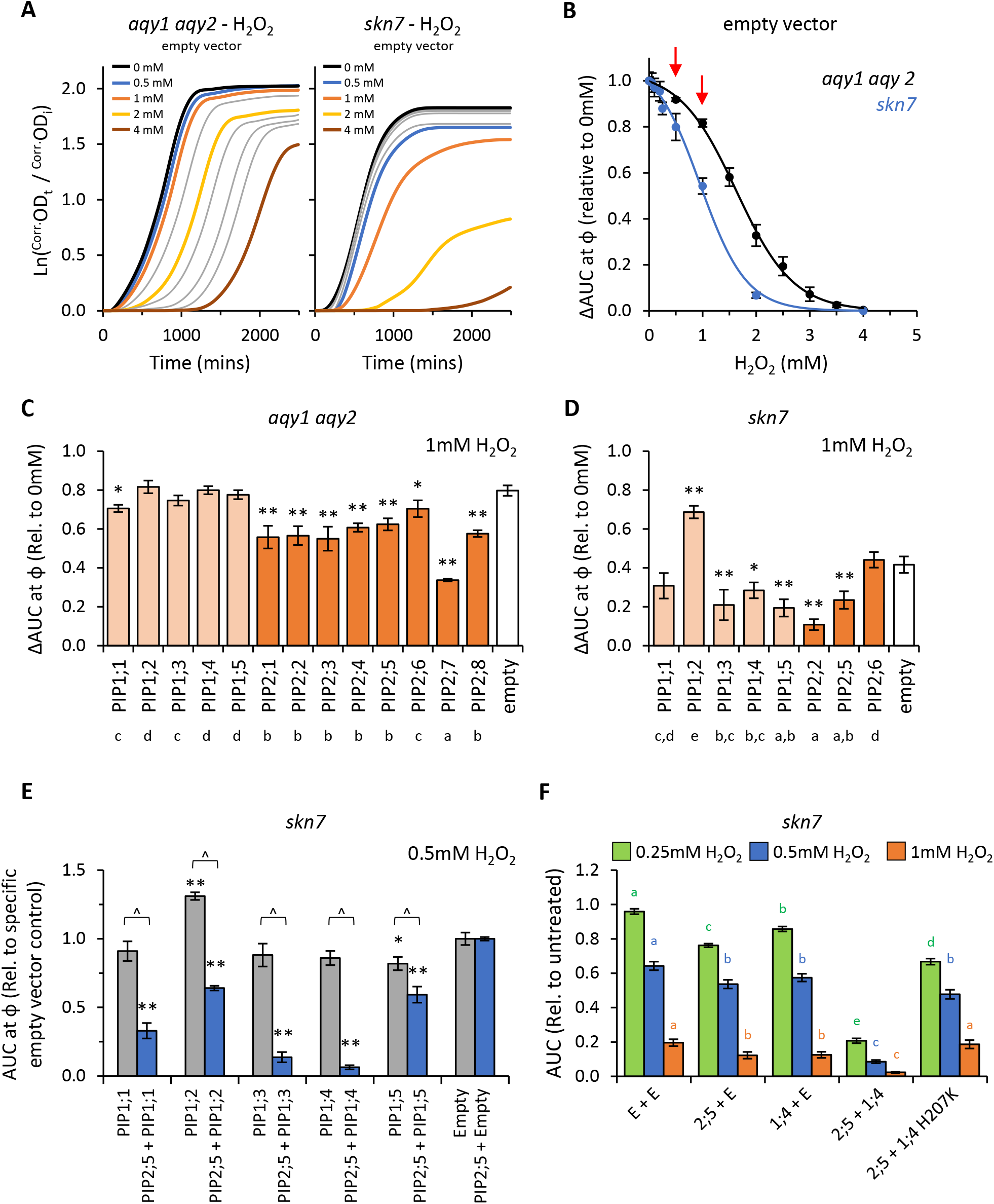
H_2_O_2_ permeability assays. **A**, Comparison of growth curves of two yeast strains, *aqy1 aqy2* or *skn7*, exposed to increasing H_2_O_2_ concentrations. **B**, Dose response curves showing relative AUC as a function of H_2_O_2_ concentration for each strain. *skn7* yeast are more sensitive to H_2_O_2_ treatment than *aqy1 aqy2* yeast. Red arrows indicate H_2_O_2_ concentrations chosen for testing yeast expressing *AtPIP*. **C**, Relative AUC for *aqy1 aqy2* yeast expressing each AtPIP gene exposed to 1mM H_2_O_2_. **D**, Relative AUC for *skn7* yeast expressing *AtPIP* genes exposed to 1mM H_2_O_2_. **E**, Relative AUC for *skn7* yeast exposed to 0.5mM H_2_O_2_ expressing *AtPIP1* singly (grey) or together with *AtPIP2*;*5* (blue). Each set is standardized to their respective empty vector control. **F**, Relative AUC for *skn7* yeast expressing various combinations of *AtPIP* genes at 0.25, 0.5 and 1mM H_2_O_2_. All error bars are SEM. For C and D, asterisks indicate statistical difference from empty vector control, ANOVA with Fishers LSD test (* *P* < 0.05; ** *P* < 0.01); letters denote different statistical rankings across both 0.5 and 1mM H_2_O_2_, ANOVA with Tukey’s test (*P* < 0.05). For E, asterisks indicate statistical difference from empty vector control, ANOVA with Fishers LSD test (* *P* < 0.05; ** *P* < 0.01); chevrons (^) indicate statistical difference between single vs. co-expression (Student’s *t* test *P* < 0.01). For F, color coded letters denote different statistical groupings within [H_2_O_2_] treatments, ANOVA with Fishers LSD test. N = 4 bio reps for B. N = 6 (2 biological reps x 3 experimental runs) for C. N = 8 across 4 experimental runs for D. For E, N = 12 across 6 experimental runs for single expressed AtPIPs and N = 6 across 3 experimental runs for co-expressed lines. N = 16 across 4 experimental runs for F.

Growth relative to the empty vector control was inhibited by 0.5mM H_2_O_2_ for all *AtPIP2* expressing *aqy1 aqy2* yeast lines except *AtPIP2*;*6* (Supplemental Fig. S6A). All AtPIP2 yeast lines grew worse than empty vector control at 1mM H_2_O_2_ (Fig. 5C), indicating that all AtPIP2 proteins can facilitate enhanced diffusion of H_2_O_2_ across the PM to some extent. AtPIP2;6 had the least and minimal implied capacity, while all other AtPIP2s were assessed as efficient H_2_O_2_ transporters, with AtPIP2;7 seemingly the most effective (Fig. 5C). The AtPIP1s showed no indication of enhancing H_2_O_2_ diffusion across the PM beyond the passive background diffusion rate represented by the empty vector *aqy1 aqy2* control, with the exception of a small effect for AtPIP1;1 at 1mM H_2_O_2_ (Fig. 5C).

When expressed in *skn7*, *AtPIP1*;*3*, *1*;*4*, and *1*;*5* conferred greater sensitivity to H_2_O_2_ (at 1mM) than empty vector control, indicating that these isoforms also facilitate H_2_O_2_ transport across the PM (Fig. 5D). The growth reduction sat between the efficient AtPIP2;2 and inefficient AtPIP2;6 H_2_O_2_ transporters originally assessed in the *aqy1 aqy2* background (Fig. 5C). Intriguingly, *skn7 AtPIP1*;*2* yeast grew consistently better than empty vector control (several independent transformation events), suggesting that expression of *AtPIP1*;*2* somehow protects *skn7* against H_2_O_2_ treatment (Fig. 5D).

Co-expression of *AtPIP2*;*5* with any of the *AtPIP1s* dramatically increased the sensitivity of *skn7* yeast to H_2_O_2_ over the *AtPIP2*;*5* + *Empty* vector control. The effect was clearly evident at 0.5mM (Fig. 5E) and even as low as 0.25mM H_2_O_2_ (Supplemental Fig. S6C), whereas 1mM H_2_O_2_ was required to observe a significant increase in *skn7* sensitivity beyond the empty vector control when the *AtPIP1s* were expressed alone (Fig. 5D; Supplemental Fig. S6B). *AtPIP2*;*5* + *AtPIP1*;*3* and *AtPIP2*;*5* + *AtPIP1*;*4 skn7* lines were the most sensitive, with ΔAUC below 15% of the *AtPIP2*;*5* + *Empty* vector control (Fig. 5E). To test whether the observed co-expressed effects related to AtPIP1 H_2_O_2_ transport or some form of hyperactivation of AtPIP2;5 H_2_O_2_ transport through hetero-oligomerization, we generated a mutant version of *AtPIP1*;*4* (*AtPIP1*;*4H207K*) with reduced channel activity (see Supplemental Materials and Methods). In an independent collection of H_2_O_2_ assays, increasing PM abundance of AtPIP1;4 through *AtPIP2*;*5* + *AtPIP1*;*4* co-expression, once again dramatically sensitized *skn7* yeast to H_2_O_2_ (Fig. 5F). However, when *AtPIP2*;*5* was co-expressed with the *AtPIP1*;*4H207K* closed gated mutant, the ΔAUC values resembled growth levels more similar to *AtPIP2*;*5* + *Empty* control (Fig. 5F). This supports the interpretation that *AtPIP1*;*4* was responsible for the enhanced H_2_O_2_ sensitivity of the *AtPIP2*;*5* + *AtPIP1*;*4* yeast. Overall, the co-expression results suggest that AtPIP1 proteins transport H_2_O_2_ more efficiently than AtPIP2 isoforms.

### Characterization of AtPIPs boric acid permeability

A range of boric acid (BA; H_3_BO_3_) concentrations were tested on *aqy1 aqy2* empty vector yeast to determine treatment doses. BA treatments mainly reduced the rate of growth (μ) (Fig. 6A; Supplemental Fig. S7). ΔAUC at ϕ relative to untreated cultures followed a single dose response curve and 20mM and 30mM BA were selected as optimal treatment concentrations (Figure 6B).

**Figure 6.**
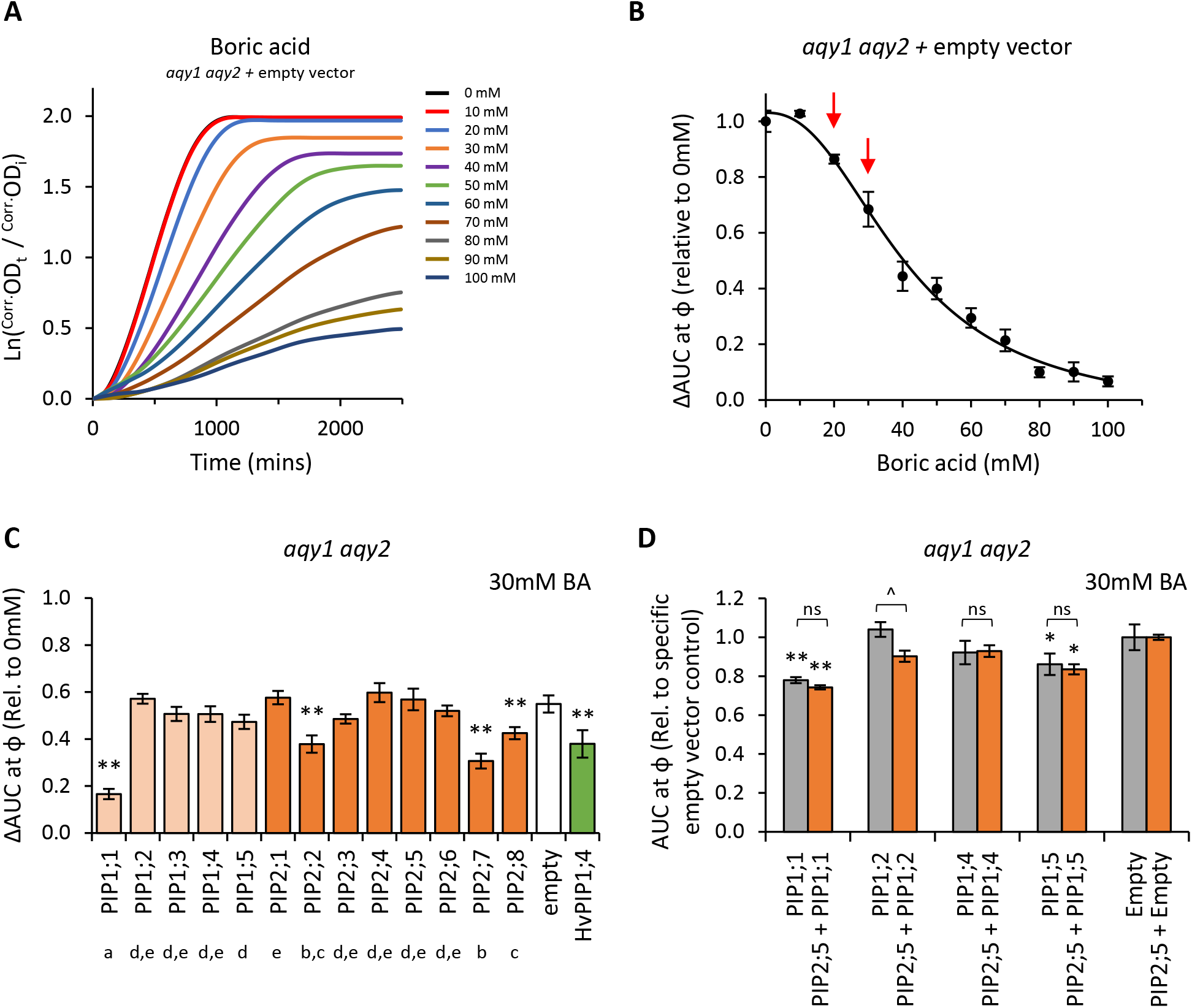
Boric acid permeability assays. **A**, Growth curves for *aqy1 aqy2* yeast exposed to increasing concentrations of boric acid (BA). **B**, Dose response curve of relative AUC as a function of boric acid concentration. Red arrows denote BA concentrations chosen for testing yeast expressing *AtPIP*. **C**, Relative AUC for *aqy1 aqy2* yeast expressing each *AtPIP* gene exposed to 30mM boric acid, with *HvPIP1*;*4* as a boric acid permeable control. **D**, Relative AUC for *aqy1 aqy2* yeast expressing *AtPIP1* singly (grey) or together with *AtPIP2*;*5* (orange) at 30mM boric acid. Each set is standardized to their respective empty vector control. All error bars are SEM. For C, asterisks indicate statistical difference from empty vector control, ANOVA with Fishers LSD test (* *P* < 0.05; ** *P* < 0.01); letters denote different statistical rankings across both 20 and 30mM boric acid, ANOVA with Tukey’s test (*P* < 0.05). For D, asterisks indicate statistical difference from respective empty vector control, ANOVA with Fishers LSD test (* *P* < 0.05; ** *P* < 0.01); chevrons (^) indicate statistical difference between single vs. co-expression (Student’s *t* test *P* < 0.01). For C and D, N = 6 across 3 experimental runs.

Five of the 13 AtPIP yeast lines were more sensitive to BA than the empty vector control (Fig. 6C, D). *AtPIP1*;*1* expressing yeast were by far the most sensitized to BA, with dramatic growth reductions even at 20mM BA. Yeast expressing *AtPIP2*;*2*, *2*;*7* and *2*;*8* had sensitivities similar to the *HvPIP1*;*4* positive control. *AtPIP1*;*5* yeast showed a small increase in BA sensitivity, which was significant in three of the four experiments (Fig. 6D; Supplemental Fig. S8A and B). Co-expression of AtPIP1s with AtPIP2;5 did not alter BA sensitivity compared to the yeast expressing *AtPIP1s* alone (Fig. 6D; Supplemental Fig. S8B).

Truncation of the cytosolic N-terminal domain of some plant AQPs, including several PIP1 isoforms, is necessary to observe boron, or similar metalloid, uptake in yeast (Bienert *et al*., 2008, Fitzpatrick and Reid, 2009, Kumar *et al*., 2014, Mosa *et al*., 2016). We generated and tested several PIP isoforms with truncations of the cytosolic N-terminal domain (*AtPIP1*;*2_Δ2-47_*, *AtPIP1*;*4_Δ2-47_* and *AtPIP1*;*5_Δ2-48_*). The truncated versions had similar sensitivity to BA as their full-length counterparts (data not shown). Overall, the results indicate that five members across both the *AtPIP1* and *AtPIP2* sub-families are capable of significant BA transport.

### Characterization of AtPIPs for urea permeability

Growth of the empty vector *ynvw1* (*dur3*) urea uptake deficient mutant was enhanced by increasing concentrations of urea; specifically through increased maximum growth rate and carrying capacity (Fig. 7A,B; Supplemental Fig. S9). None of the AtPIPs improved urea uptake, whereas the positive control *AtTIP2*;*3* (Dynowski *et al*., 2008a) clearly complemented the *dur3* phenotype at 4mM urea (Fig. 9C). With 12mM urea, all yeast lines grew similarly to the empty vector control (Fig. 9C). This indicates that firstly, all AtPIP yeast cultures were healthy and capable of growing better in response to increased urea diffusion. Secondly, the overall urea influx across the PM at 12mM exceeded the growth complementation provided by urea transport through AtTIP2;3. None of the AtPIPs were significantly permeable to urea.

**Figure 7.**
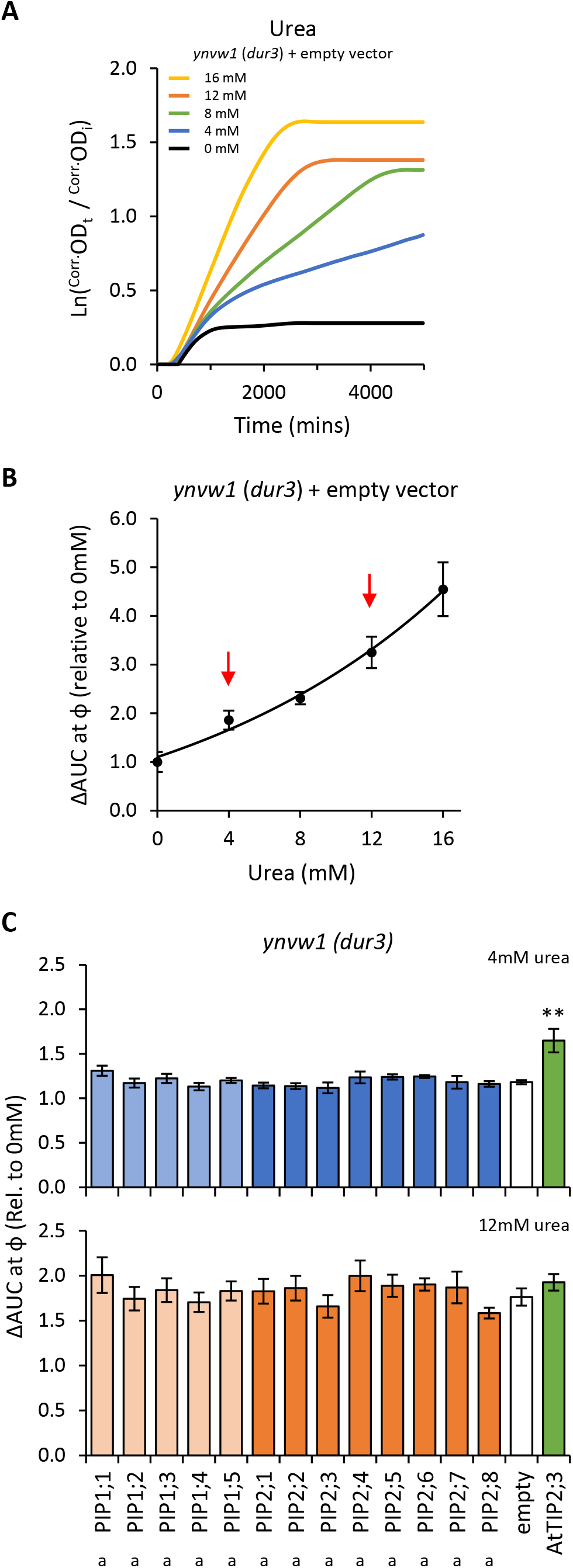
Urea permeability assays. **A**, Growth curves of *ynvw1 (dur3*) yeast supplied with increasing concentrations of urea. **B**, Relative AUC as a function of urea concentration. Red arrows denote urea concentrations chosen for testing yeast expressing *AtPIP*. **C**, Relative AUC for yeast expressing each *AtPIP* grown with 4 or 12mM urea, with *AtTIP2*;*3* as a urea permeable control. All error bars are SEM. For C, asterisks indicate statistical difference from empty vector control, ANOVA with Fishers LSD test (* *P* < 0.05; ** *P* < 0.01); letters denote statistical rankings across both 4 and 12mM urea, ANOVA with Tukey’s test (*P* < 0.05). For C, N = 6 across 3 experimental runs.

**Figure 8.**
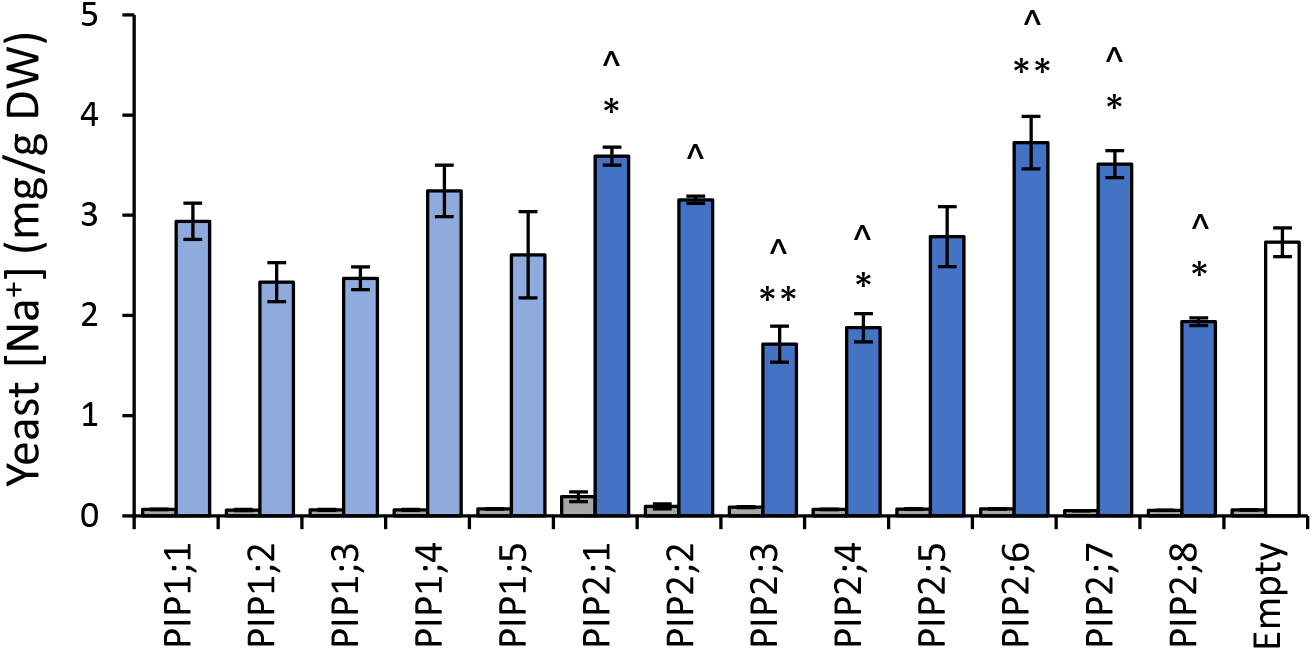
Na^+^ permeability assay. Yeast cellular sodium content before (grey) and after (blue) exposure to 70mM NaCl for 40 mins. Error bars are SEM. Asterisks indicate statistical difference from empty vector control, ANOVA with Fishers LSD test (* *P* < 0.05; ** *P* < 0.01). Chevrons (^) indicate statistical difference from empty vector control, Student’s *t* test *P* < 0.05. N = 3 for AtPIPs and N = 2 for empty vector.

**Figure 9.**
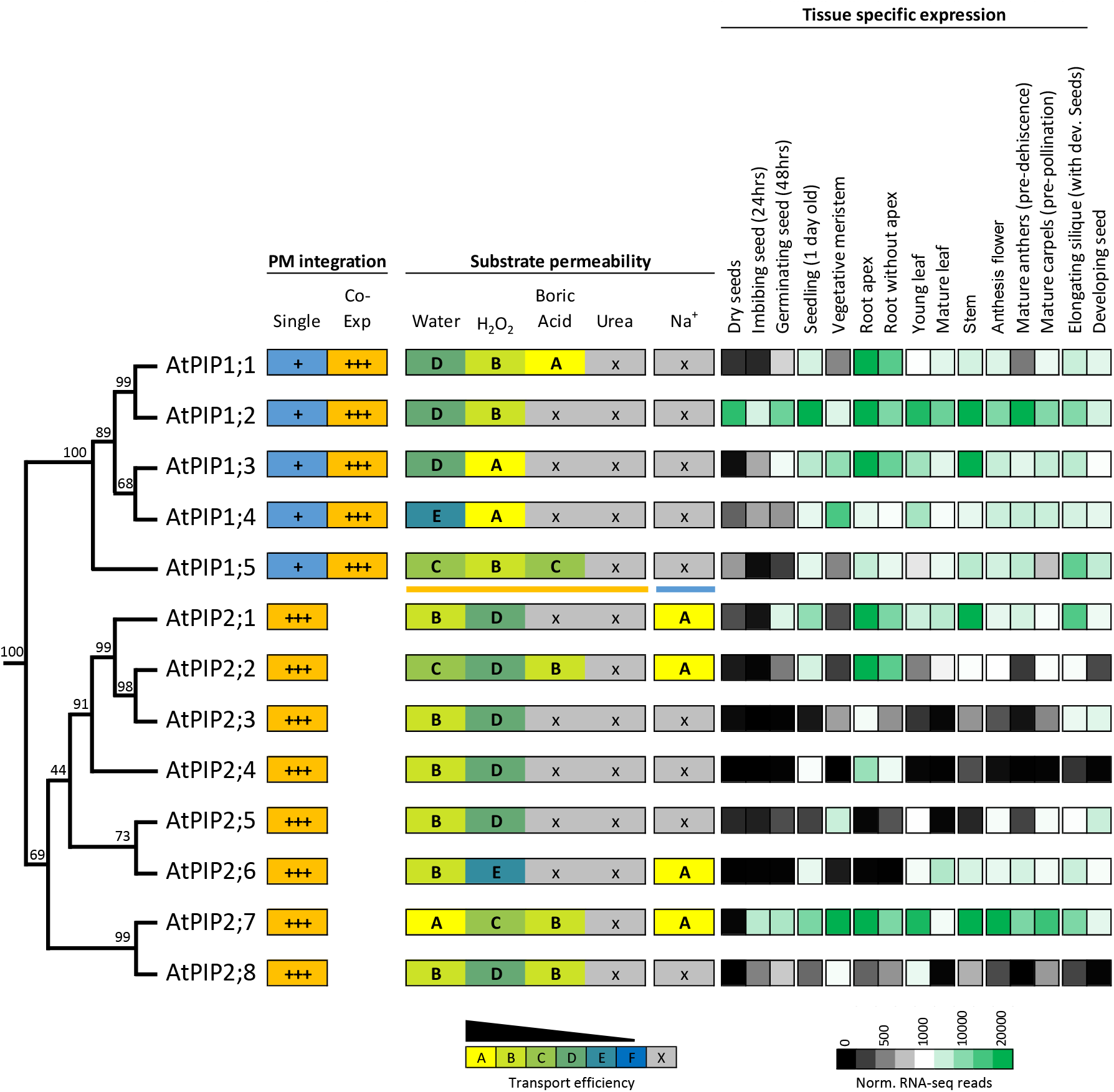
Summary of permeability and expression data for the AtPIP isoforms. The phylogenetic relationship is shown on the left, followed by strength of integration into the plasma membrane (PM) when expressed singly or co-expressed with a PIP2 (PIP1s only). Substrate permeabilities are shown in the center, and relative gene expression across different tissues during development, are shown on the right. The phylogeny is full protein sequence, using neighbor-joining method from MUSCLE alignments of protein sequences, with confidence levels (%) of branch points generated through bootstrapping analysis (n = 1000). Permeability and transport efficiencies for AtPIP1 are based on co-expression with AtPIP2;5 for water, H_2_O_2_, boric acid and urea (orange underline below AtPIP1;5) and singly expressed AtPIP1s for Na^+^ permeability (blue line under AtPIP1;5).

### Characterization of AtPIPs for Na^+^ ion permeability

To assess AtPIP potential for Na^+^ transport, we quantified Na^+^ accumulation in AtPIP expressing yeast, following NaCl treatments (Qiu *et al*., 2020). Exposing yeast to 70mM NaCl resulted in a ~40-fold increase in the Na^+^ content of *aqy1 aqy2* yeast cells relative to yeast from media without additional NaCl (Fig. 8). The five AtPIP1 isoforms and AtPIP2;5 accumulated Na^+^ similar to the empty vector control. Yeast producing AtPIP2;1, 2;2, 2;6 and 2;7 accumulated more Na^+^, while yeast producing AtPIP2;3, 2;4, and 2;8 accumulated less Na^+^ than empty vector control. AtPIP2;1 served as a positive control (Byrt *et al*., 2017).

### The evolutionary relationship, substrate profiles, and gene expression patterns of AtPIPs

Protein sequence alignments reveal the high homology between AtPIPs (Supplemental Figure S10). Motifs associated with substrate selectivity (i.e. NPA, ar/R and Froger’s positions) are essentially identical among the AtPIPs (Supplemental Table S2). Gross differences are seen in the longer N-terminal and shorter C-terminal domains of AtPIP1s compared to AtPIP2s, and variation in the length of loop A (Supplemental Table S2). Phylogenetic analysis shows that AtPIPs divide into discrete sub-clades that show distinct relationships with their substrate profiles and organ level gene expression (Fig. 9). For example, the *AtPIP1*;*1* and *1*;*2* paralogs appear to have undergone substantial functional diversification based on their gene expression patterns. *AtPIP1*;*2* is the most abundantly and constitutively expressed of all *AtPIPs*, even detected at high levels in dry seed. *AtPIP1*;*1*, is mainly expressed in roots, being ~6-fold less prevalent in aerial tissues. This diversification in expression patterns could relate to boric acid transport being present in AtPIP1;1, but absent in AtPIP1;2 (Fig. 9). The AtPIP1;3 and 1;4 paralog pair, appear to have evolved as highly efficient transporters of H_2_O_2_ while being the least efficient at water transport of the AtPIPs. Both genes are broadly expressed with largely overlapping expression domains, which together with their similar transport profiles points towards possible functional redundancy. *AtPIP1*;*3* differs from *AtPIP1*;*4* by being more highly expressed in general, especially in the root and stem. *AtPIP1*;*3* expression is also up-regulated during seed imbibition and seedling germination, whereas *AtPIP1*;*4* is only weakly expressed at this stage of development (Fig. 9). Intriguingly, AtPIP1;5 sits as a phylogenetic outgroup within the AtPIP1 clade, and transports all three substrates that AtPIP1s could transport (water, H_2_O_2_ and boric acid). AtPIP1;5 was ranked as the most efficient AtPIP1 water transporter (Fig. 9) and *AtPIP1*;*5* transcripts are particularly abundant in elongating siliques and the developing seed within.

Among the AtPIP2 isoforms, AtPIP2;7 has the most diverse substrate profile and expression patterns, being capable of transporting water, H_2_O_2_, boric acid, and Na^+^ ions at comparatively high efficiency. *AtPIP2*;*7* is expressed at high levels in most tissues, with the exception of mature leaves and dry seed, but is upregulated during seed imbibition and germination (Fig. 9). Its closest relative, AtPIP2;8, is also capable of transporting water, H_2_O_2_, and boric acid, but *AtPIP2*;*8* has relatively low expression under non-stressed growth conditions (Fig. 9). This reveals that *AtPIP2*;*8* is either highly cell specific, conditionally expressed, or that AtPIP2;7 is the dominant isoform of this closely related pair. The AtPIP2;5 and AtPIP2;6 phylogenetic pair are noteworthy as being the least efficient H_2_O_2_ transporters of all AtPIPs and they are not expressed in roots (Fig. 5C and 9). *AtPIP2*;*5* is expressed in meristematic tissue and developing seed, and *AtPIP2*;*6* expression is localized to aerial vegetative and reproductive tissues (Fig. 9).

## Discussion

### High-throughput yeast micro-cultivation assays for testing AQP substrate permeability profiles

Yeast-based systems for the heterologous expression and functional assessment of aquaporins offer numerous advantages over other systems such as oocytes, liposomes, and artificial membranes. Key advantages include: a large range of well-characterized mutant *S. cerevisiae* strains which can be used for testing different compounds; monitoring growth is simple; scalable to high-throughput processing; the power of sampling a yeast population versus single cell/event sampling in other systems.

Many studies show that altered growth in response to various chemical treatments of AQP expressing yeast reflects an enhanced intracellular accumulation of the tested substrate (Bienert *et al*., 2007, Bienert *et al*., 2008, Dynowski *et al*., 2008b, Fitzpatrick and Reid, 2009, Bienert *et al*., 2011, Kumar *et al*., 2014, Mao and Sun, 2015, To *et al*., 2015, Mosa *et al*., 2016, Rhee *et al*., 2017, Wang *et al*., 2019). We did not detect any indirect effects of AQP expression on yeast susceptibility to chemical treatments (Supplemental Note 1). Since liquid cultures provide superior substrate exposure and enable detection of smaller phenotypic changes relative to yeast grown on solid plates (Toussaint *et al*., 2006, Marešová and Sychrová, 2007, Hung *et al*., 2018), we developed a liquid micro-cultivation system enabling high-throughput, quantitative real-time monitoring of yeast growth and changes induced by treatments. The 96-well plate format offers room for multiple samples in one experiment, simplifying statistical evaluation. Optical density measurement removed the element of human subjectivity used to assess yeast spots. The implementation of a dynamic measuring point ϕ, enabled standardized evaluation between different AQP expressing yeast lines. Differential growth responses due to increased substrate diffusion into the yeast were captured by the single parameter, AUC.

High AQP contents in heterologous systems are critical for accurate assessment of functional capacity and to avoid false-negative permeabilities (Bienert *et al*., 2014). We maximized the likelihood of high AtPIP production by careful design of our AtPIP yeast expression constructs. The AtPIPs must also integrate into the yeast plasma membrane in order to affect substrate transport into the yeast cell. We found that AtPIP2s localize efficiently to the PM, while AtPIP1s co-localize to the PM and ER. Poor PM localization of PIP1s expressed alone in heterologous systems is a common phenomenon (Yaneff *et al*., 2015), likely due to sequence differences in the diacidic, LxxxA and C-terminal phosphorylation protein motifs known to control PIP2 PM trafficking (Supplemental Table 2)(Chevalier and Chaumont, 2015). The exact composition of diacidic and LxxxA motifs vary, particularly between the phylogenetically distinct [2;1, 2;2, 2;3, 2;4] and [2;5, 2;6, 2;7, 2;8] groups (Supplemental Table 2), yet all AtPIP2s localized efficiently to the yeast PM. In plants, the phylogenetically distinct AtPIP2;1 and AtPIP2;7 also localize efficiently to the PM (Sorieul *et al*., 2011, Hachez *et al*., 2014). This reveals flexibility in these motif sequences that must work together with other domains (e.g. TMH2; Wang *et al*., 2019) to control ER to PM trafficking. PIP2 proteins can physically interact with PIP1s and facilitate PM integration in both host and heterologous systems (Jozefkowicz *et al*., 2017). We enhanced AtPIP1 PM localization by co-expression with AtPIP2;5, thereby enabling the comparison of transport efficiencies.

### AtPIP water permeability

Water permeability is the most extensively studied function of PIPs across species. Most AtPIPs have been confirmed to transport water (AtPIP1;1, 1;2, 1;3, 2;1, 2;2, 2;3, 2;4, 2;6, and 2;7) (Kammerloher *et al*., 1994, Tournaire-Roux *et al*., 2003, Heckwolf *et al*., 2011, Byrt *et al*., 2017, Kourghi *et al*., 2017, Wang *et al*., 2020a). These assessments are from different studies and systems making it difficult to directly compare transport efficiencies. Here, water permeability was assessed for the complete set of AtPIPs using a freeze-thaw assay that we established for rapidly evaluating water transport capacity of AQPs. We found that all AtPIP isoforms transport water, with AtPIP2s more efficient than AtPIP1s. Studies that concluded PIP1s have low/no permeability to water, may reflect the inefficient targeting of PIP1s to the PM in heterologous systems [reviewed in (Yaneff *et al*., 2015)].

PIPs provide a transcellular route for water flow in the plant, from water uptake by roots to transpiration loss from aerial tissues (Groszmann *et al*., 2017). Both AtPIP1 and AtPIP2 isoforms play major roles in water flow in Arabidopsis (Javot *et al*., 2003, Prado *et al*., 2013, Sade *et al*., 2014). Overlapping expression patterns suggest substantial functional redundancy, which limits the ability of reverse genetic studies to resolve the contribution of each AtPIP to water flow. For example, single loss-of-function mutants of high leaf-expressing isoforms *Atpip1*;*2*, *Atpip2*;*1* and *Atpip2*;*6* each show a ~20% reduction in rosette hydraulic conductivity, which worsens to ~39% in the triple mutant (Prado *et al*., 2013). Our observations that *AtPIP2*;*7* is highly permeable to water and is abundantly expressed in developing leaves (Figure 9), suggests it may also contribute to rosette hydraulic conductivity. Similarly, redundancy for root hydraulic conductance is likely given that the 10-20% reductions seen in single *Atpip* mutants falls short of the ~64% decrease achieved using AQP chemical blockers (Maurel *et al*., 2015). Four of the seven *AtPIPs* abundantly expressed in roots (Figure 9), are the more water permeable AtPIP2 isoforms (AtPIP2;1, 2;2, 2;4, 2;7) and thus strong candidates for multiple knock-out mutant studies.

More intricate developmental processes relying on cell-to-cell water movement through AtPIPs are emerging. For example, guard cell closure (Grondin *et al*., 2015), lateral root emergence (Péret *et al*., 2012), and pollen germination on stigmatic papillae (Windari *et al*., 2021). A number of AtPIPs are expressed in the flower, developing silique and seeds. In these tissues, AtPIP water transport could have roles in petal expansion, anther/pollen development, and assist in the supply of nutrients to the developing seed as seen in other species (Hoai *et al*., 2020, Wang *et al*., 2020b).

### AtPIP H_2_O_2_ permeability

All AtPIPs are capable of transporting H_2_O_2_ when expressed in yeast (Figure 5), which is consistent with the similar physicochemical properties of H_2_O_2_ and water (Almasalmeh *et al*., 2014). Previous growth-based assessments with yeast did not assign H_2_O_2_ permeability to AtPIP1 isoforms and showed mixed results for AtPIP2 isoforms (Hooijmaijers *et al*., 2012, Wang *et al*., 2019, Wang *et al*., 2020a). This may have been due to inadequate protein production, insufficient PM targeting, choice of yeast strain, sub-optimal H_2_O_2_ concentrations, or use of solid medium spot growth assays.

The potential for H_2_O_2_ transport through AtPIP1s was recently hinted at using AtPIP1/2 chimeric proteins that more effectively localize to the PM (Wang *et al*., 2019). However, in addition to harboring PM targeting motifs, the substituted PIP2 domains also contribute to the pore lining, making it uncertain how representative these chimeric proteins are of native AtPIP1 function. In our system, we were able to show that native AtPIP1 proteins are indeed capable of transporting H_2_O_2_, and when efficiently targeted to the PM through co-expression, are potentially more effective transporters of H_2_O_2_ than AtPIP2 isoforms.

H_2_O_2_ is an indispensable signaling molecule involved in many aspects of plant growth, biotic defense and abiotic stress responses, reliant on AQPs to facilitate its movement between sub-cellular compartments and cells (Černý *et al*., 2018, Fichman *et al*., 2021). The diversity of AtPIP expression patterns and H_2_O_2_ transport efficiencies, enable fine tuning of H_2_O_2_ signaling. Direct physiological evidence in Arabidopsis is emerging, with H_2_O_2_ transport through AtPIP2;1 involved in triggering stomatal closure (Rodrigues *et al*., 2017) and mediating systemic acquired acclimation to abiotic stress (Fichman *et al*., 2021), and AtPIP1;4 mediating H_2_O_2_ triggered immunity against pathogen attack (Tian *et al*., 2016). Our results show that the *AtPIP1*;*3*/*1*;*4* paralogs have evolved into highly efficient H_2_O_2_ transporters with largely overlapping tissue-specific expression patterns. This redundancy suggests that *AtPIP1*;*3* could also mediate H_2_O_2_ signaling for plant immunity. Supporting this idea, H_2_O_2_ translocation into the cell is decreased but not eliminated in the *atpip1*;*4* single mutant (Tian *et al*., 2016); and only *AtPIP1*;*4* and *AtPIP1*;*3* are rapidly up-regulated in response to H_2_O_2_ treatment of leaves (Hooijmaijers *et al*., 2012). The latter would be a consistent response to the apoplastic H_2_O_2_ produced upon pathogen recognition and facilitating its entry into the cell to trigger immune responses (Tian *et al*., 2016). *AtPIP1*;*3* transcripts are not present in dry seed, but are substantially induced during seed imbibition and germination. Hydrating seed releases H_2_O_2_ as a signal to promote germination, and may involve AtPIP1;3, which would be consistent with the involvement of AQPs in the germination process (Hoai *et al*., 2020). Further investigation into a role for AtPIP1;3 in plant immunity and seed germination appears warranted.

### AtPIP boric acid permeability

Five AtPIPs were permeable to boric acid, with a ranking of AtPIP1;1 > AtPIP2;2 = AtPIP2;7 = AtPIP2;8 > AtPIP1;5. Boric acid permeability is generally associated with NIP-type AQPs (Pommerrenig *et al*., 2015). However, a growing number of PIP isoforms from different species are being found capable of transporting boron in heterologous systems; ZmPIP1;1 (Dordas *et al*., 2000), OsPIP1;3 and OsPIP2;6 (Mosa *et al*., 2016), OsPIP2;4 and OsPIP2;7 (Kumar *et al*., 2014), and HvPIP1;3 and HvPIP1;4 (Fitzpatrick and Reid, 2009). A native physiological role for PIP boron transport is not yet confirmed in any species, but improved tolerance to boron toxicity in Arabidopsis over-expressing boron permeable rice PIPs, points towards a possible role (Kumar *et al*., 2014, Mosa *et al*., 2016).

Boron permeable AtPIPs are expressed in all tissue types and may help coordinate uptake and distribution of this essential micronutrient, and provide tolerance via efflux under toxic concentrations. AtPIP1;1 was an efficient boron transporter, but not its paralog AtPIP1;2. *AtPIP1*;*1* expression is unaltered in roots and minimally in shoots under toxic boron conditions, whereas *AtPIP1*;*2* is substantially repressed (Macho-Rivero *et al*., 2018). *AtPIP1*;*1* which is permeable to boron, is predominantly expressed in roots and differentially expressed in response to boron. This suggests it has undergone substantial functional diversification since duplication with *AtPIP1;2. AtPIP1*;*2* is widely and highly expressed and facilitates CO_2_ diffusion into chloroplasts for photosynthesis (Heckwolf *et al*., 2011), whereas we suggest AtPIP1;1 may be specialized for micronutrient uptake from the soil.

### AtPIP urea permeability

Urea differs massively from water with respect to size, polarity and other physicochemical properties. No AtPIP was capable of permeating urea, which is consistent with urea being too large to pass through the narrow aperture of the AtPIP a/R filter (Supplemental Table S2)(Dynowski *et al*., 2008a, Dynowski *et al*., 2008b).

### AtPIP Na^+^ permeability

Yeast tolerance of salt toxicity is associated with osmo-resistance (Stratford *et al*., 2019), meaning that AtPIP water transport could confound growth data for AtPIP expressing yeast grown at high salt concentrations. Therefore, assessment of AtPIP Na^+^ permeability from yeast growth requires a tailored mutant (Sychrova, 2004). Instead, to screen for AtPIP Na^+^ permeability, we quantified intracellular yeast Na^+^ content directly. We confirmed previous reports of Na^+^ permeability for AtPIP2;1 and AtPIP2;2 (Byrt *et al*., 2017, Qiu *et al*., 2020), and observed that AtPIP2;6 and AtPIP2;7 also appear permeable to Na^+^. The latter is at odds with previous electrophysiological experiments on *AtPIP2*;*7* expressing oocytes that report AtPIP2;7 is not permeable to Na^+^ (Kourghi *et al*., 2017). The contrasting findings could reflect different heterologous expression systems and detection techniques, but investigation of post-translational regulation of AtPIP2;7 function is warranted.

We observed no enhanced Na^+^ accumulation in yeast expressing AtPIP1s alone. Since the central pore, formed in the middle of tetrameric AQP complex, is the pathway for monovalent ions (Yool and Weinstein, 2002), we did not screen yeast co-expressing AtPIP1s with AtPIP2;5. This would change the structure of the central pore and make interpretation of results ambiguous, as seen for CO_2_ and Na^+^ transport through the central pore of PIP hetero-tetramers (Otto *et al*., 2010, Byrt *et al*., 2017).

The dual permeability to water and solutes of certain AtPIPs may help build high turgor during cell expansion. For example, AtPIP2;1 is involved in lateral root emergence where the primordia pushes through the overlying tissues (Péret *et al*., 2012). Our observations that AtPIP2;7 has dual water and solute transport capacity and is upregulated during seed imbibition and germination, implies a role aiding the massive influx of water needed for the radicle to puncture through the seed coat. Moreover, expression of *AtPIP2*;*7* in seeds responds to two antagonistically acting phytohormones (GA and ABA) that regulate seed dormancy versus germination (Hoai *et al*., 2020).

### Why the differences in efficiency between isoforms?

We observed differences among AtPIPs in their efficiency to transport water and H_2_O_2_ and capability to permeate boric acid or Na^+^. This is puzzling given the near identical residue signatures of motifs classically considered to govern substrate selectivity (i.e. NPA, ar/R, and Froger’s positions) (Supplemental Table S2; Figure S10), and indicates the involvement of other domains yet to be defined. Variation in transport efficiency for water and H_2_O_2_ is likely to be associated with subtle differences in residues forming the monomeric pore that alter the number of hydrogen bonds with the substrate, or that shift, even slightly, the spatial configuration of the pore diameter (Horner *et al*., 2015, Mom *et al*., 2021). Differences in the sensitivity of gating regulation and the degree of ‘openness’ or ‘open probability’ is another possible factor (Kourghi *et al*., 2017, Vitali *et al*., 2019, Qiu *et al*., 2020). Na^+^ transport was only detected for some AtPIPs, pointing to differences in central pore features (Yool and Weinstein, 2002). The route for boric acid through PIPs is unknown, but mutant analysis suggests the monomeric pore is most likely (Dynowski *et al*., 2008b). However, we cannot exclude the central pore given its hydrophobic profile and hypothesized ability to open wider through helix rotation (Tyerman *et al*., 2021). Structural changes to the central pore of hetero-tetramers would also account for the inability to improve AtPIP1;1 and AtPIP1;5 boric acid permeability when co-expressed with AtPIP2;5.

The limited sequence differences between the AtPIPs (Supplemental Figure S10), should make identification of substrate specificity residues easier and feasible to explore through mutation approaches.

## Conclusion

Using a micro-volume yeast-based system we developed comparative substrate permeability profiles for the entire AtPIP subfamily. The validity of our micro-volume yeast system was assured by bench-marking against published AtPIP permeability data from different systems. Comparison between AtPIP isoforms and across multiple substrates allowed for more informative conclusions. For example, although AtPIP2;6 was permeable to water, it was an inefficient H_2_O_2_ transporter.

Our substrate profiles align with known biological roles of AtPIPs and will help uncover further physiological roles obscured by genetic redundancy. The rich resources in Arabidopsis (mutants, expression data, physiological studies etc.) should allow evaluation of permeability profiles and reveal physiological significance more readily than in other species.

Transgenic manipulation of AQPs to improve yield or stress tolerance in various plant species has mixed outcomes (Chaumont and Tyerman, 2017). Neutral or negative phenotypes could be related to off-target substrate transport through the manipulated AQP, or insufficient transport efficiency to yield a desirable effect. Broader comparative profiling would provide a vital strategic tool for selecting ‘better’ candidates towards fit-for-purpose translational AQP applications. For example, using AtPIP2;4 over AtPIP2;7 for more exclusive water permeability, or using AtPIP1;3 if highly efficient H_2_O_2_ transport is required over water.

Microplate readers suitable for AQP yeast assays are becoming readily affordable, which should favour the use of liquid cultures over solid medium for growth evaluations. Our system can be expanded to other AQP types. Building a catalogue of transport capacity from a large number of AQPs will help clarify biological roles and decipher the nuanced characteristics of transport selectivity and efficiency necessary for future engineering of AQPs for specific biotechnological applications.

## Materials and methods

Detailed material and methods are provided as Supplemental Information. Briefly, *AtPIP* and control gene coding sequences were commercially synthesised (Genscript) as gateway-enabled entry constructs and cloned into destination vectors from the Advanced Gateway^®^ series of yeast expression plasmids (Alberti *et al*., 2007) to create the various yeast expression clones. These were transformed into appropriate yeast strains using Frozen-EZ yeast Transformation Kit II (Zymo Research). *AtPIP-GFP* were used to evaluate heterologous AtPIP production, with GFP signal detected in concentrated yeast cultures using the Infinite M1000 Pro plate reader (TECAN). Subcellular localization in yeast cells was performed using confocal microscopy on a Zeiss LSM780 confocal laser-scanning microscope (Carl Zeiss) operated by Zen Black software. Quantification of AtPIP2;5 interactions with AtPIP1 proteins using the Y2H mbSUS was performed as per (Grefen *et al*., 2007). Yeast spheroplasts were generated using zymolyase digestion (Zymo Research) and spheroplast bursting due to osmotic shock measured using a Cary 60 UV-VIS (Agilent) spectrophotometer with OD650 reading at 0.1 sec intervals. Micro-volume yeast cultures were cultivated and OD readings measured using a Spectrostar Nano microplate reader (BMG, Germany) in Nunc-96 400 μL flat bottom untreated 96-well plates (Thermo Scientific Cat#243656) with lid and 200μl culture volume per well. Default cycling conditions for yeast growth assays were: 250 cycles at 10 mins per cycle (total time ~42-50 hrs); incubated at 30°C with a slightly warmer lid; shaking frequency of 400 rpm in double orbital shaking mode; 5 mins shaking per cycle prior to the OD reading, with the remaining time the plate sitting idle on the incubation plate; OD readings invoke orbital averaging at scan diameter of 4mm and 22 flashes per well, recording at 650nm. OD_650_ readings minus the blank were corrected for non-linearity using our pre-determined calibration function to generate ^Corr.^OD_650_ at a 1cm path-length, the data was converted into growth curves that were smoothed using several filters and finally log (LN) transformed using ^Corr^OD_650_ at time ‘t’ divided by the initial starting OD (^Corr.^OD_i_). Specifics of freeze-thaw, H_2_O_2_, boric acid, urea, and NaCl treatments are detailed in Supplemental Materials and Methods.

## List of author contributions

MG conceived the original screening, framework, and research plans and made the yeast expressing the AtPIP constructs; MG and ADR developed the micro-cultivation methodology and established optimal treatment concentrations; MG developed data processing methodology, MG performed the AtPIP yeast screening experiments and analysis; MG and WC performed AtPIP interaction and yeast spheroplast analysis; JQ and SAM developed and performed the sodium uptake assay with supervision by CSB; MG, JRE, CSB and ADR analyzed the data and wrote the article. All authors critically reviewed the manuscript. MG agrees to serve as the author responsible for contact and ensures communication.

## Acknowledgements

The authors acknowledge the facilities and the scientific and technical assistance of Darryl Webb of Microscopy Australia at the Advanced Imaging Precinct at the Australian National University; a facility funded by the ANU, and State and Federal Governments of Australia. We also thank Peter Dahl of the S. Hohmann lab, for providing us the 10560-6B wild type and *aqy1 aqy2* mutant yeast strains. Gerd P. Bienert for supplying the *ynvw1* yeast mutant. Christopher Grefen for components of the Y2H mbSUS.

## Supplemental Data

**Supplemental Figure S1.** Adjusting for non-linearity of OD measurements at high cell density.

**Supplemental Figure S2.** The highly active *GPD* promoter confers greater AQP enhanced water permeability over the less active *TPI1* promoter.

**Supplemental Figure S3.** Quantification of AtPIP protein abundance in intact yeast.

**Supplemental Figure S4.** Establishing the freeze-thaw assay for water permeability.

**Supplemental Figure S5.** Calibrating H_2_O_2_ treatments for yeast growth assay.

**Supplemental Figure S6.** H_2_O_2_ permeability assays.

**Supplemental Figure S7.** Calibrating boric acid treatments for yeast growth assay.

**Supplemental Figure S8.** Boric acid permeability assays.

**Supplemental Figure S9.** Calibrating urea treatments for yeast growth assay.

**Supplemental Figure S10.** AtPIP family protein sequence alignment.

**Supplemental Figure S11.** Correlation analysis examining AtPIP induced changes in inherent yeast growth characteristics and possible indirect effects on response to treatments.

**Supplemental Figure S12.** Growth curve processing.

**Supplemental Figure S13.** Deriving μ, *λ*, and κ, from a processed growth curve.

**Supplemental Table S1.** *AtPIP* codon compatibility for heterologous expression in yeast and growth characteristics of *AtPIP* expressing yeast lines.

**Supplemental Table S2.** Protein domain lengths and amino acid composition of AtPIPs at known substrate selectivity positions and other important motifs.

