## Supplemental Information for "Permeability profiling of all 13 Arabidopsis PIP aquaporins using a high throughput yeast approach"

#### Supplemental Note 1

**AtPIP effects on yeast growth are specific to increased membrane permeability**

Heterologous expression of AtPIPs can alter the inherent growth characteristics of the host yeast strain independent of substrate treatment, with some AtPIPs affecting lag phase (λ), growth rate (μ), and carrying capacity (κ) more than others (Supplemental Table S1). These changes may influence the sensitivity of the yeast to treatment and confound our conclusions. However, no correlations were observed between treated ΔAUC values when compared against values for λ, μ, and κ from untreated cultures (Supplemental Figure S11). Therefore, the differential growth responses of certain *AtPIP* yeast lines to specific treatments appears the direct result of AtPIP enhancement of membrane permeability to the tested substrate and not due to indirect effects through subtle changes to inherent growth characteristics.

#### Supplemental Materials and Methods

**Yeast strain genotypes**

Wild type strain: 10560-6B (WT AQY1 AQY2; background Σ1278b; genotype: Mat α; leu2::hisG; trp1::hisG, his3::hisG; ura352). Provided by Peter Dahl of the S. Hohmann lab.

*aqy1 aqy2*: YKF 1771 (null aqy1 aqy2; background Σ1278b; genotype: Mat α; leu2::hisG; trp1::hisG, his3::hisG; ura352 aqy1D::KanMX aqy2D::KanMX). Provided by Peter Dahl of the S. Hohmann lab.

*skn7*: ATCC® Number: 4002900™ (null skn7; background BY4741 genotype: Mat a his3Δ1 leu2Δ0 met15Δ0 ura3Δ0 ΔSKN7). Obtained from ATCC (www.atcc.org).

*ynvw1* (*dur3*): (null dur3; background Σ23346c; genotype: Mat a, Δura3, Δdur3). Provided by Patrick Bienert of the Nicolaus von Wirén lab.

**Cloning of *AtPIP* genes into yeast expression vectors**

*AtPIP* coding sequences were commercially synthesised (Genscript) as gateway-enabled entry vectors (i.e. included flanking attL sites). A second set of *AtPIP* genes without the stop codon were also made, for use in GFP C-terminal fusion constructs. The *AtPIP* CoDing Sequences (CDS) were cloned into plasmids from the Advanced Gateway® adapted pRS series of yeast expression plasmids (Alberti *et al.*, 2007) using Gateway *LR Clonase II* enzyme mix (Invitrogen™). The library was obtained through Addgene (https://www.addgene.org/)(kit #1000000011). pAG423GPD-ccdB (MG0515; HIS) or pAG426GPD-ccdB (MG0517; URA) were used for full-length CDS clones for permeability assays. pAG426GPD-ccdB-EGFP (MG0528; URA) was used for GFP fusions (i.e. AtPIP CDS without stop codon). All E.coli cloning steps used One Shot™ OmniMAX™ 2 T1R Chemically Competent E. coli cells (Initrogen). All final plasmids were sanger sequenced to confirm accuracy of the clones using; Wizard®

Plus SV Minipreps DNA Purification Systems (Promega), BigDye® sequencing chemistry (Thermofisher Scientific), and ZR DNA Sequencing Clean-Up Kit (Zymo Research). Final expression vectors were transformed in yeast using the Frozen-EZ yeast Transformation Kit II (Zymo Research, Los Angeles, USA). MG0515 constructs were transformed into *Δaqy1aqy2* and *Δskn7* yeast strains and used for freeze-thaw, $H_2O_2$ and boric acid assays. MG0517 constructs were transformed into *Δdur3* (*Δynvw1*) cells and used for the urea assay.

**Cloning of control AQPs for different substrate assays**

A yeast codon optimized version of barley *PIP1;4* (*HvPIP1;4* was chosen as a positive control, as it has been reported to transport boric acid in yeast from agar spot and cellular boron content assessments (Fitzpatrick and Reid, 2009). The codon usage of *HvPIP1;4* was adapted for expression in the yeast genome (see CAI adaptation below), which is especially important for expressing monocot derived genes in yeast (Bienert *et al.*, 2014). The native coding sequence of *AtTIP2;3* (AT5G47450) was cloned and used as a positive control for urea uptake assay (Dynowski *et al.*, 2008a). Both *HvPIP1;4* and *AtTIP2;3* were cloned like the *AtPIP*s, including modification to Kozak sequences (see below). We generated and used *AtROP6* (AT4G35020) as an additional plasma membrane-localized protein control in the Y2H mbSUS. *AtKAT1* (AT5G46240) was provided as part of the Y2H mbSUS kit gifted to us by Christopher Grefen (University of Tuebingen).

**Design considerations of yeast expression constructs**

High AQP protein production in the yeast is essential for robust functional evaluations and to minimize potential false negative assessments of permeability (Bienert *et al.*, 2014). To assure ample protein production, we considered several regulatory components in designing our primary *AtPIP* yeast transgenes.

***Gene copy number***

Gene copy number is an important component for obtaining high gene and subsequently protein expression in yeast (Chen *et al.*, 2012). We used the Advanced Gateway® adapted pRS series of episomal yeast expression vectors (Alberti *et al.*, 2007), to enable high-throughput cloning and to allow for multiple copies of the *AQP* harboring plasmids per yeast cell. The plasmid library provided a choice of origin of replication, promoter, and auxotrophic selection marker. We chose the 'high-copy' 2μ origin of replication to elevate plasmid copy number, which should be bolstered by our use of the GPD promoter, yielding in the range of 20-30 plasmid copies per cell (Karim *et al.*, 2013). Only plasmids with either the histidine (HIS) [MG0515; pAG423GPD-ccdB] or uracil (URA) [MG0517; pAG426GPD-ccdB; URA selection] selection markers were used, as these both appear to have equally minimal plasmid load burdens on yeast growth in selection medium (Karim *et al.*, 2013).

***Promoter***

The choice of the right promoter is a crucial point for efficient gene expression in yeast. The *APQ* transgenes were all driven by the highly active promoter of the yeast *GLYCERALDEHYDE-3-PHOSPHATE DEHYDROGENASE (GAPDH) ISOZYME 3* gene (*GPD*); also referred to as *TRIOSE-PHOSPHATE DEHYDROGENASE* (*TDH3*). Of the most commonly used yeast promoters, *GPD* (*TDH3*) is among the strongest drivers of transgene expression in the glucose-rich medium of our micro-cultivation conditions (Partow *et al.*, 2010, Da Silva and Srikrishnan, 2012, Lee *et al.*, 2013, Peng *et al.*, 2015). The benefits of using a stronger promoter was evident in our initial trials where enhancements in AtPIP2;3 driven water permeability were substantially greater when using the strong GPD promoter over the less active *TRIOSE PHOSPHATE ISOMERASE* (*TPI1*) promoter (Supplemental Fig. S2).

***Terminator***

There was only a single choice of terminator in the Advanced Gateway® yeast vector kit, with all constructs utilizing the well-known *CYC1* yeast terminator. Although alternative terminators may help yield higher protein levels with different promoters, these appear to provide little additional benefit when using the highly active *GPD* promoter (Curran *et al.*, 2013).

***Codon compatibility***

Beyond transcriptional determinants, high AQP heterologous protein production in the yeast will rely on the efficiency of translating the mRNA. This is partially dependent on the congruency of codon usage between the foreign *AQP* sequence and the yeast genome, and can be measured by the Codon Adaptation Index (Sharp and Li, 1987). The CAI values for all 13 *AtPIPs* against yeast codon preferences range between 0.68 - 0.74 with a GC content of 49% - 52% (Supplemental Table S1). These values are similar to their native CAI values within Arabidopsis (0.73 - 0.81) and also within the range of the native yeast *AQP* genes (*AQY1*: 0.71 CAI and 51% G+C; *AQY2*: 0.79 CAI and 46% G+C). Given this level of harmonization, we chose to clone the native *AtPIP* sequences into the yeast expression vectors.

***Kozak sequence***

Translation efficiency is also influenced by the Kozak sequence surrounding the AUG of the mRNA that triggers the ribosome to commence translation (Kozak, 2005). The 5'UTR sequences of all *AtPIP* transgenes were designed identical and included aspects of a Kozak sequence favoring higher protein expression in yeast (Supplemental Table S1) (Dvir *et al.*, 2013, Li *et al.*, 2017). Downstream of the AUG was dictated by the native *AtPIP* sequences and were generally acceptable sequence configurations (Supplemental Table S1). Most *AtPIP*s have a guanine at position +4 (i.e. AUGG), considered quintessential for higher protein production in eukaryotes, widely used by yeast genes, and correlated with greater human AQP protein production in yeast (Kozak, 2005, Öberg *et al.*, 2009, Hernández *et al.*, 2019) (Supplemental Table S1). *AtPIP2;5* to *AtPIP2;8*, have an adenine or thymine at position +4 (Supplemental Table S1), generally related to more moderate protein production in yeast, but do have the favorable cytosine at +5 (Robbins-Pianka *et al.*, 2010).

**Codon Adaptation Index value calculations**

CAI values were calculated using the CAI calculation tool through the CAIcal Server: http://genomes.urv.es/CAIcal/ (Puigbò *et al.*, 2008). Codon usage tables were obtained through the Kazusa database: http://www.kazusa.or.jp/codon/ (Nakamura *et al.*, 2000). CAI values for each of the AtPIPs against the Arabidopsis and *S. cerevisiae* genomes are presented in (Supplemental Table S1).

**Protein quantification and subcellular localizations of heterologous AtPIP proteins in yeast**

C-terminal GFP fusions are an effective means of evaluating heterologous protein production in cultures of viable growing intact yeast, correlating well with traditional Westerns, and for monitoring subcellular localizations of membrane bound proteins (Albano *et al.*, 1998, Ghaemmaghami *et al.*, 2003, Drew *et al.*, 2006, Drew *et al.*, 2008, Scharff-Poulsen and Pedersen, 2013). We generated a collection of *AtPIP-GFP* transgenes using the pAG426GPD-ccdB-EGFP (MG0528) vector from the Advanced Gateway® adapted pRS series of episomal yeast expression vectors (Alberti *et al.*, 2007) that, besides the C-terminal GFP fusion, are identical in design to our primary transgenes used for the yeast permeability assays. Additionally, as the AtPIPs are of similar size to each other (278-291 aa; Supplemental Table S1) and larger than eGFP (239 aa), the production of the AtPIP-GFP chimeric protein should primarily reflect the translation efficiencies of the different AtPIPs. As such, the *AtPIP-GFP* constructs are a suitable proxy for evaluating heterologous AtPIP production and subcellular localization of our primary *AtPIP* constructs.

***Quantification of AtPIP protein abundance in intact yeast***

A single ~5 mm diameter colony of each *AtPIP-GFP* expressing yeast line being tested was placed in 3ml yeast nitrogen base (YNB) with 2% Glucose (v/v) and supplemented with appropriate amino drop-out (DO) (i.e. -URA), and cultured at 30°C at 230rpm for 24hrs. 50ul of overnight culture was added to 450ul of YNB medium and measured at $OD_{660}$ (OD reading should fall within the predetermined linear portion of your spectrophotometer). YNB SD DO medium was added to overnight cultures to adjust OD to a $^{Corr.}OD_{660}$ = 5. 2.5ml of adjusted culture was transferred to a fresh tube and centrifuge for 5mins at 3000g. The supernatant was poured off and the pellet resuspended in 220ul of appropriate YNB SD DO medium. The concentrated culture was now in the range of $^{Corr.}OD_{660}$ = 60. For each construct, 3 wells of a black 96-well plate were used (Nunc™ F96 MicroWell Black nontreated. Thermoscientific Cat#237105). 200ul of the yeast concentrate was place in well 1, and 100ul of YNB SD DO medium was added to wells 2 and 3. Doubling dilutions were performed using 100ul from well 1 to well 2 and then to well 3, giving concentrations of 100%, 50% and 25%. Just prior to measurement, cells were resuspended using a multichannel pipette. GFP fluorescence was measured using an Infinite M1000 Pro plate reader (TECAN) with Gain 115, Z-position 19300, excitation wavelength 488nm, emission detection 512nm, excitation and emission bandwidth 5nm, 50 Flashes, at Flash frequency of 400 Hz, Integration time 20µs, and Settle time 50ms. GFP fluorescence readings over the 3 concentrations (i.e. 100%, 50%, 25% cell density) needed to be consistent and in the linear detection range ($R^2 \geq 0.99$). Cell concentration of the 100% well was confirmed by repeated measurements of 5ul of 100% cell concentrate to 495ul fresh YNB SD medium dilutions at $OD_{660}$. Yeast not expressing GFP were also measured to determine (minimal) background fluorescence levels. Protein abundance was reported as AtPIP-GFP fluorescence values minus background fluorescence, standardized to cell $OD_{660}$ = 1. Six independent replicate experiments were performed for each *AtPIP-GFP* expression yeast line.

***AtPIP-GFP subcellular localization in yeast***

Single ~5 mm diameter colonies of *AtPIP-GFP* expressing *Δaqy1aqy2* yeast were used to inoculate 1ml starter culture (YNB DO with 2% Glucose v/v) and grown to stationary phase (24 hrs, 30 °C, 230 rpm). The over-night cultures were diluted 4-fold and incubated for a further 4hrs prior to GFP visualisation. 2µl of settled yeast cells were spotted on a polylysine coated slide (Thermofisher scientific) and mounted with 22x22mm cover slip and sealed with nail varnish. Sub-cellular GFP signal was visualised on a Zeiss LSM780 confocal laser-scanning microscope (Carl Zeiss) operated by Zen Black software and a DIC x40 oil immersion lens. eGFP was excited at 488nm and emission was recorded at 495-570 nm with 1.35 AU pinhole and master and digital gains identically set for all images and analysis. The SEC63-RFP ER marker was obtained through Addgene (pSM1959; Addgene plasmid #41837; donated by Susan Michaelis; (Metzger *et al.*, 2008)). RFP was excited at 561 nm and emission was captured at 565-735 nm with 1.10 AU pinhole with a quick scan time to avoid bleaching of RFP. Over 30 cells for each of the AtPIP-GFP lines were scored to access membrane localisation tendencies and performed across two replicate experiments, with localisation patterns between the genotypes consistent across sessions. We also performed subcellular localisation analysis in the *Δskn7* background, which concluded similar findings.

#### Micro-volume yeast growth system

***Adjusting for non-linearity of OD measurements at high cell density***

Obtaining accurate OD values of batch cultures at high cell densities ideally requires aliquots to be diluted to within the linear detection range (i.e. where OD ≈ cell concentration). This is simply not feasible in a high-throughput set-up. Instead, the non-linear OD relationship can be compensated for by comparing ‘recorded’ ODs against ‘true’ ODs calculated from dilution factors (Supplemental Figure 1). Fitting a regression function will allow for the relationship between ‘recorded’ and ‘true’ ODs to be readily calculated (i.e. ‘recorded’ $OD_{650}$ at 200ul into ‘true’ $OD_{650}$ at 1cm pathlength). Such a calibration function needs to be established independently for a given machine (or at least different brand machines), as the pathlength, detector, light beam angle, aperture etc, will vary. Calibration can also be performed between two different spectrophotometers to correct/collate data collected between different machines. We used the Spectrostar Nano microplate reader (BMG Labtech,

Germany) for this study, which is an affordable basic model plate reader capable of shaking, incubations, and multiple point reading, run by a versatile software program and complete with its own dedicated database (MARS – BMG Labtech).

***Shaking modes***

Linear, orbital and double orbital shaken modes were trialed for their capacity to agitate cultures and keep yeast in suspension. Regardless of the mode and shaking speed (up to 1000rpm), yeast eventually sedimented. We found that the dispersal of sedimentation on the bottom of the well, was markedly important for the reproducibility of growth curves. Linear (left-right) and orbital (circular) shaking were not suitable, as they produced cell clumping as ridges (linear) or a ring shape (orbital) with distinct regions absent of cells. Double orbital (figure eight) shaking produced and maintained a homogenous lawn of yeast cell sediment across the base of the well throughout the experiment, and is the only shaking mode we recommend. Shaking mode and frequency is also imperative for the continued mixing of the medium and exposure of cells to the supplemented agents being tested. A shaking velocity of 400rpm was chosen for its ability to quickly and thoroughly disperse small droplets of dye gently loaded into wells containing 200ul of yeast growth medium. A shaking duration of 5 mins prior to the OD reading (occurring usually every 10 mins) was deemed a suitable balance between sufficient mixing and considerations of wear and tear on equipment.

***OD reading mode***

'Orbital averaging' mode on the Spectrostar Nano microplate reader (BMG, Germany) was invoked for OD readings. In this mode, the plate reader collected data from a 4mm diameter orbit (22 read points) and reported the average value for a more stable representative OD reading with reduced deviation between replicate wells. A 650nM wavelength was chosen for OD readings as our medium and yeast generally do not absorb strongly at this longer wavelength at any time throughout the growth cycle. This means that OD readings will be dominated mainly by the single factor of scattering that will generally provide a more consistent representation of cell concentration.

***Incubation***

Growth assays were incubated at the optimal growth temperature of 30$^{o}$C for yeast. The Spectrostar Nano microplate reader (BMG, Germany) heats the lid of the 96-well plate slightly warmer to avoid condensation during runs. Although the entire stage is heated, we invoked the software option to shift the plate back to the full heating plate during the initial idle incubation period of each cycle, to avoid the potential cool spot directly above the optical sensor.

Our standard incubation periods were for ~42hrs, except for urea, where growth was monitored for ~50hrs. This is because although yeast can use urea as a sole source of nitrogen, growth rates are slower than when supplied with a preferred nitrogen sources such as ammonium (Godard *et al.*, 2007).

***Default kinetic conditions for yeast growth assay***

The default cycling conditions used for our yeast growth assays were: 250 cycles at 10mins per cycle (total time ~42 hrs)*; incubated at 30$^{o}$C with a slightly warmer lid to avoid condensation; shaking frequency of 400 rpm; double orbital shaking mode; 5 mins shaking per cycle prior to the OD reading, with the remaining time the plate sitting idle on the incubation plate and away from the colder optical reading region; OD readings invoke orbital averaging at scan diameter of 4mm and 22 flashes per well, recording at 650nM. * 12 min cycles for urea supplementation assay (total time ~50 hrs).

***Micro-culture volume and starting cell concentration***

We settled on 200ul culture volumes. 100ul cultures appeared to mix insufficiently, with yeast occasionally clumping and producing erratic growth curves. 300ul was excessive, with shaking exceeding the retention capacity of the well. Cultures with higher initial cell concentrations that produced a lawn of cells provided the most reproducible growth curves. Too low of an initial cell density, produced independent scattered colonies and inconsistent growth curves often with erratic curve anomalies (e.g. sharp peaks or sudden drops). Our cultures began with an initial $OD_{650}$ of ~0.4 as measured in the 200ul volume within the 96-well plate. Ideally, the initial cell density provides a discernable lag phase for untreated cultures, but still allows sufficient growth of treated cultures within a reasonable time period.

***Processing of growth curves***

Reliable yeast population growth characteristics (e.g. λ, μ, and κ) were obtained from growth curves built from frequent OD data acquisition and critical post-processing steps. Changes in growth characteristics revealed how treatments affected yeast growth and allowed quantitative comparison between successive yeast cultures. The implementation of a dynamic measuring point ϕ, allowed for standardized evaluation between different AQP expressing yeast lines, regardless of any reasonable impact that the AQP may exert on inherent growth of untreated yeast. Differential growth responses due to increased substrate diffusion into the yeast were captured by the single parameter, AUC.

Yeast growth data was processed using a custom-built fit-for-purpose pipeline in Microsoft Excel that draws on existing algorithms to process and calculate characteristics of yeast population growth curves (Warringer and Blomberg, 2003, Toussaint *et al.*, 2006, Olsen *et al.*, 2010, Hall *et al.*, 2014, Jung *et al.*, 2015, Fernandez-Ricaud *et al.*, 2016). Most importantly, our pipeline allows for visualization of each processing step and comparisons between treatments, for full scrutiny of obtained curves and AUC measurements. The pipeline consists of; **Removing of background:** In each experimental run, several wells are dedicated 'blanks' consisting of non-inoculate growth medium complete with appropriate chemical treatment additives. 'Blank' well values were removed from the corresponding yeast growth OD values at each time-point during the growth cycle. **OD calibration:** The OD calibration function was then applied (see: Adjusting for non-linearity of OD measurements at high cell density), providing $^{Corr.}OD_{650}$ values (Figure 1A). **Adjusting for settling time:** During the first 60 - 180 mins, there is a sharp increase in OD values (Supplementary Figure 12A). This mostly occurs due to an increase in cell size as the yeast exit from the starved $G_0$ state and reestablish a size permissive for replication. The OD values then settle to a 'norm', representing the actual OD of the lag-phase period. The OD values during the settling period are adjusted commensurate to the acclimated lag-phase OD (Supplementary Figure 12B). **Curve smoothing:** Several filters were applied to reduce noise across the collective OD readings of a growth curve. (i) Median filter - Considers 5 consecutive data points and replaces the middle value with the median of all 5. Median filtering minimises the impact of aberrant spike readings impacting later smoothing process and growth characteristic extractions (Supplementary Figure 12C). The first and last two data points of the entire series are cloned (i.e. extending the data beyond the recorded timepoints) to avoid edge cropping when applying this filter at the ends of the data collected. (ii) Mean filter - A sliding window of 3 consecutive data points, where the middle value in each window is replaced by the mean of the series. It is a light smoothing operation that does not distort the true data trend (Supplementary Figure 12D). (iii) Removing negative slopes filter - Sweeps each data series in a step-wise manner replacing any OD value lower than its predecessor with the predecessor itself (i.e. removing negative slopes between consecutive points) (Supplementary Figure 12E). This filter is used to compensate for occasional minor stochastic changes in OD readings brought about by, for example, bubble formation that drifts in and out of the light beam. (iv) Cubic scanning filter - As a final smoothing step, a cubic smoothing spline is applied, which is highly flexible and has useful smoothing properties that are not bound by assumed population curve behaviours. A sliding window consisting of 1 central focal data point and 30 flanking points either sides, are fitted with a cubic equation ($Ax^3+Bx^2+Cx+D$) and the central focal value adjusted to fit the regression line (Supplementary Figure 12F). Prior to the application of this filter, the first and last data points are cloned 30 times to prevent edge cropping when the calculation window nears the start or end of the data series. The application of these four filters provides a smoothed growth curve from which subsequent growth characteristic parameters can be more accurately derived (Supplemental Figure 12G). **Log transformation:** The smoothed growth data is then log transformed (LN) and plotted as Ln($^{Corr\cdot}OD_t$ / $^{Corr\cdot}OD_i$), where $^{Corr\cdot}OD_i$ is the initial OD (Supplementary Figure 12H). $^{Corr\cdot}OD_t$ / $^{Corr\cdot}OD_i$ provides a relative assessment, easier computation of AUC values, helps correct minor variation between reps, and can be applied when ODi are generally equivalent (which should be the case if pre-assay dilutions of samples was properly conducted). Some treatments alter the ODi, for example the lysis of cells occurring as a result of freeze-thaw treatments (Supplementary Figure 12J). The reduced ODi artificially inflates the ODt/ODi ratio compared to control and can distort comparisons of growth curves. Therefore, when reporting in Ln($^{Corr\cdot}OD_t$ / $^{Corr\cdot}OD_i$) mode, ODi is derived from the untreated control (or highest essential nutrient treatment) and applied to all cultures of the given series (Supplemental Figure 12J). Our pipeline enabled toggling between LN ($^{Corr\cdot}$OD) and LN($^{Corr\cdot}OD_t$ / $^{Corr\cdot}OD_i$) to explore treatment effects and decide on the most appropriate output format.

***Extraction of key growth characteristics***

The exponential growth phase was determined using linear slopes consisting of a seven time point sliding window (Focal time-point +/- 3 flanking time points) (red data values in Supplemental Figure S13). The point of maximum slope was identified and a flanking 10% deviation was used to define the extremes of a maximum exponential range (blue data values in Supplemental Figure S13). A linear regression line was applied to this region and used to derive maximum growth rate (μ). Lag phase (λ) was defined as the intersect point of the maximum exponential slope and the X-axis (orange in Supplemental Figure S13). Stationary phase, or carrying capacity (κ), was determined as the point the population growth rate hits virtually zero with the measuring point (ϕ) occurring just prior at the point population growth rate falls to 5% of the maximum slope (green in Supplemental Figure S13). AUC was determined using a segmented trapezoid approach. For each time interval up to the measuring point, segments of AUC was calculated using Area = $(LN(OD/ODi)_n + LN\ (OD/ODi)_{n+1})/2 \times (t_2\text{-}t_1)$, which were then totalled to provide AUC.

#### Treatments of AQP expressing yeast lines

##### Freeze-thaw assay

Yeast are sensitive to very rapid freezing events, with the formation of intracellular ice crystals causing cell damage and death. Yeast freeze-thaw survivorship involves various factors, but is mainly associated with how rapidly water can efflux from the cell across the PM (Cabrera *et al.*, 2020), which in turn is correlated with the abundance of water permeable AQPs (Tanghe *et al.*, 2002, Tanghe *et al.*, 2004, Soveral *et al.*, 2006, Will *et al.*, 2010). Therefore, water permeability of AQP variants can be rapidly determined in yeast by screening for improved freeze-thaw survivorship (To *et al.*, 2015).

The common practice was for several independent yeast transformants, once identified on selection plates, to be immediately cultured, cells harvested, and stored at on glycerol at -80°C. Each independent yeast transformant was considered a biological replicate. 1ml cultures (in 14ml round bottom tubes), were grown at 30°C, shaking at 250rpm, until stationary phase for each AQP expressing line being tested. 7.5ul of each culture was spotted onto appropriate agar-based DO selection medium and incubated for 2-3 days. One-third of a spot was used to inoculate an assay starter culture consisting of 1.25ml appropriate YNB SD DO medium, at 30°C, shaking at 250rpm for 24 hours (~$OD_{650}$ 1). An aliquot of the starter culture was diluted in 1ml at $6\times10^6$ cells/mL in YNB SD DO medium in a 2ml microfuge tube and incubated at 30°C for 60 mins. 250μL of each culture was aliquoted to an appropriate number of 1.5ml microfuge tubes; one tube for each round of freeze-thaw (FT) treatment being applied. The zero FT tubes were placed on ice (untreated control), while the others were progressed through (generally two) freeze-thaw cycles, and placed on ice once they reached the desired treatments. Each FT cycle consisted of the yeast culture being rapidly frozen in liquid nitrogen for 30 seconds, and thawed in a 30°C water bath for 20 mins. For each construct, 200ul of 'Untreated' and 'Treated' yeast were transferred from the 1.5ml microfuge tubes into the 96-well plate for micro-cultivation and growth detection.

**$H_2O_2$ assay**

The reactive oxygen species (ROS) hydrogen peroxide ($H_2O_2$), can impair yeast growth when it exceeds the protective mechanisms of the cell (Jamieson, 1998). A range of $H_2O_2$ concentrations added to the external growth medium were tested to determine ideal treatment doses. *AtPIP2;1, 2;4* and *2;5* served as controls having been shown to transport $H_2O_2$ in yeast using both agar spot growth and low-throughput fluorescent-dye-based cellular accumulation assays (Dynowski *et al.*, 2008b, Mao and Sun, 2015, Wang *et al.*, 2019). We also used the *skn7* null mutant yeast strain, which is compromised in its antioxidant buffering capacity and is therefore more sensitive to $H_2O_2$ treatment (Krems *et al.*, 1996, Morgan *et al.*, 1997).

To check that co-expression of *AtPIP1* was not associated with some form of hyperactivation of the AtPIP2;5 through hetero-oligomerization, we made a mutant version of *AtPIP1;4* (*AtPIP1;4H207K*) with hindered channel activity but retained hetero-oligomerization ability. The histidine at position 207 in AtPIP1;4 represents a strongly conserved residue located in a cytosolic loop of PIP proteins that is involved in gating of the monomeric pore (Törnroth-Horsefield *et al.*, 2006). Mutation to a positively charged Lysine(K) mimics histidine protonation, which favors pore closure and reduces transport of substrates, including $H_2O_2$ (Tournaire-Roux *et al.*, 2003, Verdoucq *et al.*, 2008, Bienert *et al.*, 2014).

*aqy1 aqy2 or skn7 y*east expressing AtPIPs or carrying the Empty vector, were grown and diluted to $6x10^6$ cells/mL as per the Freeze-thaw assay (above). 200µL microcultures of each AtPIP/Empty vector were distributed in 96-well plate comprised of 190µL of yeast and 10µL $H_2O_2$ to achieved desired final treatment concentrations (with water for the 0mM $H_2O_2$ treatment).

**Boric acid assay**

Boron is essential for yeast growth, but at high concentrations is toxic. At moderate concentrations (< 80mM) it acts as a fungistatic agent, slowing down proliferation but not killing the cell (Bennett *et al.*, 1999, Schmidt *et al.*, 2010). A range of boric acid (BA; $H_3BO_3$) concentrations were tested against *aqy1 aqy2* empty vector yeast. Barley *PIP1;4* (*HvPIP1;4*) was used as a positive control, having been reported to transport BA in yeast from agar spot and cellular boron content assessments (Fitzpatrick and Reid, 2009). We codon optimized *HvPIP1;4* to favor high expression in yeast given that monocot AQP sequences express poorly in yeast (Bienert *et al.*, 2014). The *HvPIP1;4* construct was designed with the same construct principles as the AtPIPs.

*aqy1 aqy2 y*east expressing AtPIPs or carrying the Empty vector were grown and diluted to $6x10^6$ cells/mL as usual. 200µL microcultures of each AtPIP/Empty vector were distributed in 96-well plate comprised of 180µL of yeast and 20µL boric acid to achieved desired final treatment concentrations (with water for the 0mM boric acid treatment).

**Urea assay**

We employed the use of the urea uptake-defective strain *ynvw1* (*dur3*), which possesses a deletion of *DUR3* urea transporter, and consequently shows compromised growth in medium containing low urea concentrations as the sole nitrogen source (Liu *et al.*, 2003). *ynvw1* yeast expressing *AtTIP2;3* (Dynowski *et al.*, 2008a) was used as a positive control.

*ynvw1* yeast spots grown on YNB plates were scraped and resuspended in 1.25mL of Yeast Basic media (YB, culture medium without nitrogen source) with 2% Glucose. Yeast cultures were diluted to $1.2x10^7$ cells/mL. 200µL microcultures for each AtPIP/Empty vector construct were distributed in 96-

well plates with 190μL of yeast and 10μL urea treatment (with water being used for the 0mM urea treatment).

**NaCl assay**

$Na^+$ uptake was quantified by spectrophotometry of lysed cell content from yeast exposed to NaCl treatment as previously described (Qiu *et al.*, 2020) except for the following modifications: the full-length of AtPIPs were sub-cloned into MG0515 (pAG423GPD-ccdB) yeast expression destination vector driven by *GDP* (*TDH3*) promoter through LR reaction. Confirmed constructs were transformed into *Saccharomyces cerevisiae aqy1 aqy2* double mutant yeast strain using Frozen-EZ Yeast Transformation II kit (Zymo Research). The yeast culture media for growing and for performing $Na^+$ uptake assays contained 2% glucose (w/v) rather than galactose.

#### Yeast spheroplast assays

The traditional yeast spheroplast bursting method for detection of AQP water permeability (Daniels *et al.*, 2006), involves placing yeast spheroplasts (i.e. yeast without a cell wall) in hypotonic solution which causes water to diffuse into the cell, leading to cellular expansion and rupture, with cell lysis monitored by a loss in optical density. Spheroplast expressing water permeable AQPs rupture more readily than control cells, with the effect proportional to their capacity to transport water (Pettersson *et al.*, 2006, Azad *et al.*, 2009). Briefly, a 5ml culture was grown to mid to late exponential phase. 2 x 2ml aliquots were centrifuged at 1500 x g room temperature for 2 mins and supernatant removed. The pellet was resuspended in 1ml of water, centrifuge at 1500 x g and the water discarded. The pellets were combined by resuspension in 850ul of freshly made 'softening medium' (100mM Hepes-KOH, pH 9.4; 10mM DTT), incubated for 30 mins at room temperature, centrifuge at 3000 x g for 4 mins at room temperature and supernatant removed. The cells were resuspended in 850ul of 'spheroplasting medium' (1xYNB, 2% glucose, 1x amino acid, 50mM Hepes-KOH pH 7.2, 1M sorbitol), centrifuged at 1500 x g for 1 min at RT, supernatant removed, and resuspended once again in 850ul of spheroplasting medium. 400ul of cells were diluted to $OD_{650}$ 3, in spheroplasting medium, 2 µl of zymolyase 100T lytic enzyme (Zymo Research) added, and incubated for 4 hrs at 30°C with gentle inversion. The kinetics of 20 µl of spheroplasts mixed with 480 µl spheroplast bursting medium (15% spheroplasting medium and water v/v) was measured using Cary 60 UV-VIS (Agilent) spectrophotometer with $OD_{650}$ reading at 0.1 sec intervals to determine if sufficient digestion was achieved. Actual assays were performed using 75 µl of spheroplasts for more consistent readings.

#### Y2H mbSUS

We used the Yeast-two-Hybrid mating-based Split-Ubiquitin System for detection of membrane bound protein-protein interactions (Y2H mbSUS) with assays preformed as described in (Grefen *et al.*, 2007). We used the ONPG assays to detect β-Gal activity, indicative of interaction strength. We screened an AtPIP interactome library that we produced using the mbSUS vectors (Groszmann, unpublished). We generated and used *AtROP6* (AT4G35020) as an additional plasma membrane-localized protein control in the Y2H mbSUS. *AtKAT1* (AT5G46240) control was provided as part of the Y2H mbSUS kit, which was gifted to us by Christopher Grefen (University of Tuebingen).

#### Statistics

All statistical comparisons (e.g. ANOVA, Ttest, etc) and curve fitting applications during calibrations of the various treatments (Supplemental Figures S4, S5, S7, and S9) were performed using Origin Pro 2020 (ver. 9.7.0.188).

#### Bioinformatics

AtPIP protein sequences were obtained through The Arabidopsis Information Resource (TAIR) (www.arabidopsis.org). Protein sequences were imported into Geneious (ver. 11.1.5) and aligned using MUSCLE sequence aligner (Edgar, 2004). Transmembrane and loop domains were defined using

TOPCONS (Tsirigos *et al.*, 2015). The AtPIP phylogeny was generated using neighbor-joining method from MUSCLE alignments of full protein sequences, with confidence levels (%) of branch points generated through bootstrapping analysis (n = 1000) using MEGA 10 (Kumar *et al.*, 2018). Normalized tissue-specific RNA-seq data was obtained through TRAVA (http://travadb.org/) (Klepikova *et al.*, 2016) and heatmaps produced using MS Excel.

### Supplemental Figures

#### Permeability profiling of all 13 Arabidopsis PIP aquaporin genes using a high throughput yeast approach

Michael Groszmann[1*], Annamaria De Rosa[1], Weihua Chen[1], Jiaen Qiu[2], Samantha A McGaughey[1], Caitlin S. Byrt[1] and John R Evans[1]

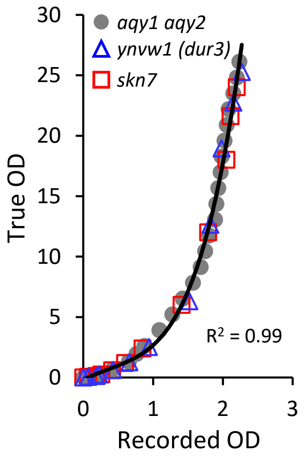


**Supplemental Figure S1. Adjusting for non-linearity of OD measurements at high cell density.** Correlative analysis between 'recorded' OD on plate reader versus 'true' OD as determined through dilution factors. The micro-volume data was fitted with a polynomial function ($y = 1.9481x^4 - 4.2474x^3 + 5.0329x^2 + 0.3441x$), that converts 'recorded' $OD_{650}$ at 200ul readings to 'true' $OD_{650}$ readings at 1cm pathlength.

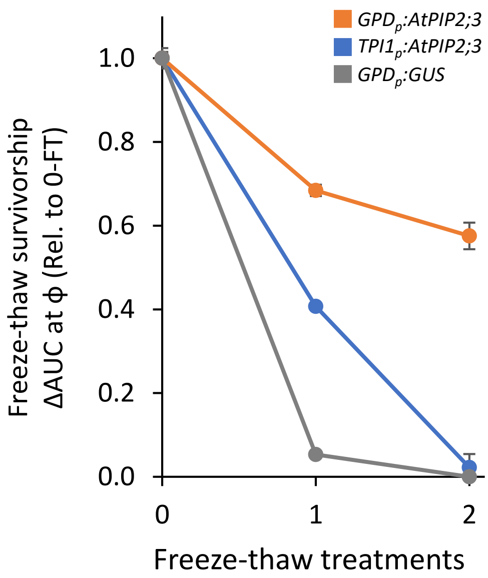


**Supplemental Figure S2. The highly active *GPD* promoter confers greater AQP enhanced water permeability over the less active *TPI1* promoter.** *aqy1 aqy2* double mutant yeast carrying $GPD_p$:AtPIP2;3 show superior rates of freeze-thaw survivorship (a proxy for water permeability) over yeast carrying $TPI_p$:AtPIP2;3. The *GPD* (*TDH3*) promoter is reported to be approximately twice as active as the *TPI1* promoter in the glucose-rich conditions of our micro-cultures (Partow *et al.*, 2010, Peng *et al.*, 2015). This substantial difference in phenotype, highlights not only the benefit of using the strong GPD promoter, but also the importance of having high AQP production to ensure robust phenotypes for evaluation. Freeze-thaw survivorship was determined by the cumulative growth (AUC) from culture inception to time-point ϕ between untreated and freeze-thawed cells. Yeast expressing the β-glucuronidase reporter gene driven by the GPD promoter was used as negative *AQP* control. Mean ± SEM, n = 3 biological replicates. *GPD - GLYCERALDEHYDE-3-PHOSPHATE DEHYDROGENASE (GAPDH) ISOZYME 3. TPI - TRIOSE PHOSPHATE ISOMERASE*

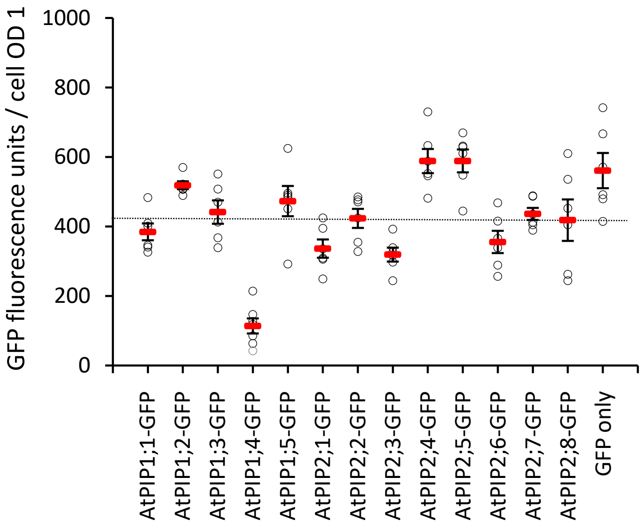


**Supplemental Figure S3. Quantification of AtPIP protein abundance in intact yeast.** GFP fluorescence intensity of yeast cultures expressing AtPIP:GFP fusion proteins were used to assess heterologous AtPIP protein production between the various lines. Assay was performed in the *aqy1 aqy2* null mutant yeast strain. Dotted line is the average GFP intensity signal across all AtPIPs. GFP fluorescence intensity standardised to a cell $^{Corr.}OD_{660} = 1$. N = 6 biological replicates across 6 experimental runs, each measured in duplicate across three dilutions (see supplemental materials and methods).

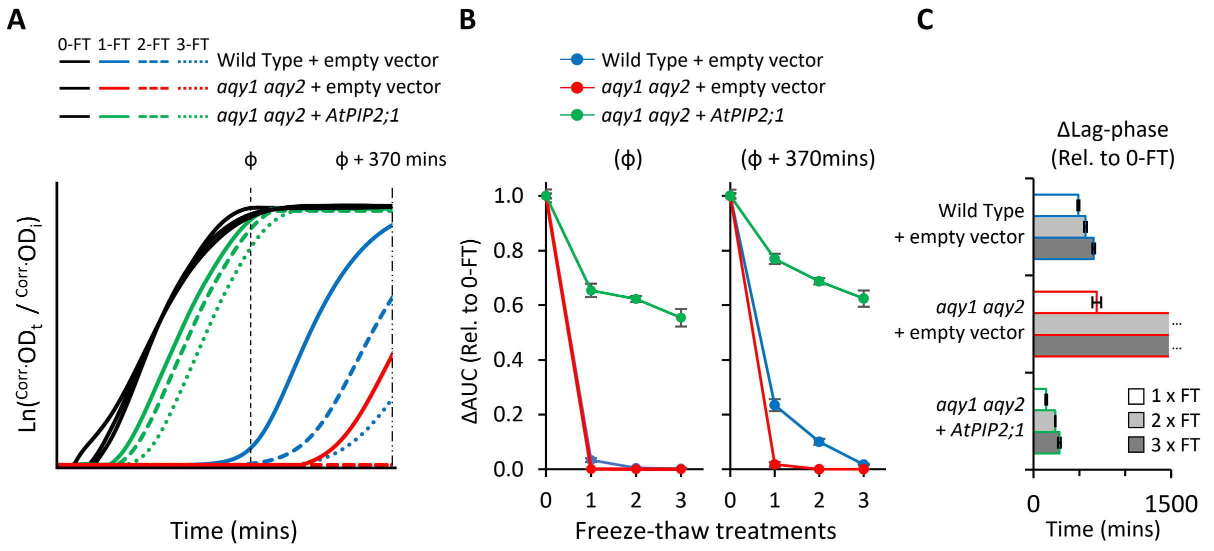


**Supplemental Figure S4. Establishing the freeze-thaw assay for water permeability. A**, Growth curves of WT, *aqy1 aqy2*, and *aqy1 aqy2* expressing *AtPIP2;1* yeast after 0, 1, 2, or 3 freeze-thaw treatments. **B**, ΔAUC values at two different measuring points, ϕ and ϕ + 370 mins, demonstrating that the *aqy1 aqy2* yeast are more sensitive than the WT yeast to freeze-thaw treatment with an incremental decrease in ΔAUC after successive freeze-thaw treatments. Expression of the water permeable *AtPIP2;1* in the *aqy1 aqy2* yeast confers a significantly higher survival rate. **C**, Freeze-thawing prolongs the lag phase, which is consistent with a reduction in the viable cell count of the starting population due to the freeze-thaw treatments.

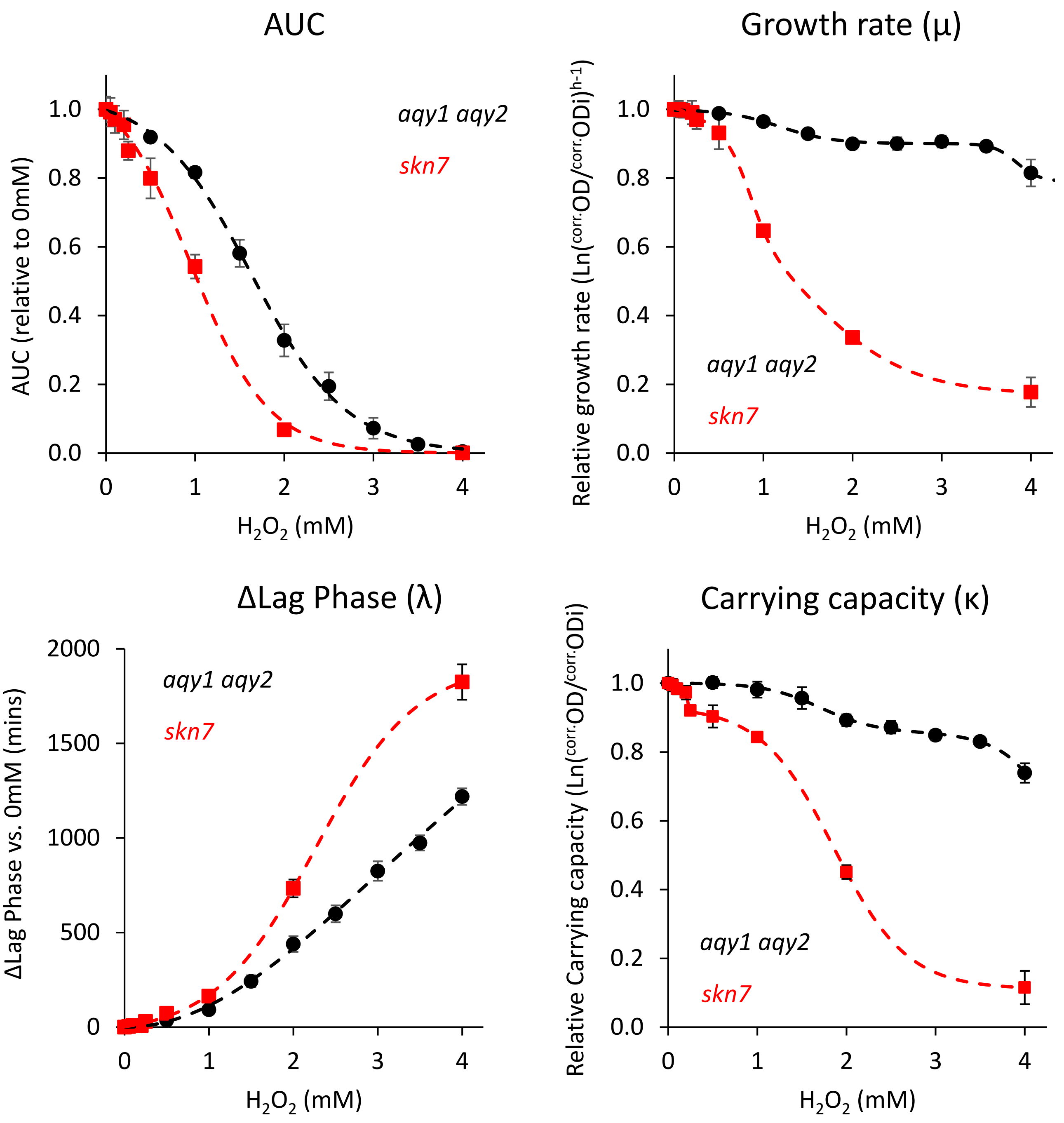


Dose Response curve: $$y = A1 + \frac{A2 - A1}{1 + 10^{(LOGx0 - x)p}}$$

Bi-Dose Response curve: $$y = A1 + (A2 - A1)\left[\frac{p}{1 + 10^{(LOGx01 - x)h1}} + \frac{1 - p}{1 + 10^{(LOGx02 - x)h2}}\right]$$

**Supplemental Figure S5. Calibrating $H_2O_2$ treatments for our yeast growth assay.** Growth responses of *aqy1 aqy2* and *skn7* yeast carrying empty vector control, to increasing concentrations of $H_2O_2$ added to the growth medium. As expected, the *skn7* mutant yeast strain, which is compromised in aspects of its antioxidant buffering capacity, was substantially more sensitive to $H_2O_2$ treatment than *aqy1 aqy2*. Both yeast strains showed phasic responses in μ and κ, indicating points where $H_2O_2$ concentrations overwhelm ROS buffering mechanisms. The growth responses captured by AUC and were fitted by a single dose response curve. Curve fitting of individual traits: μ - Bi-Dose response curve; λ - Dose response curve; κ - Bi-Dose response curve.

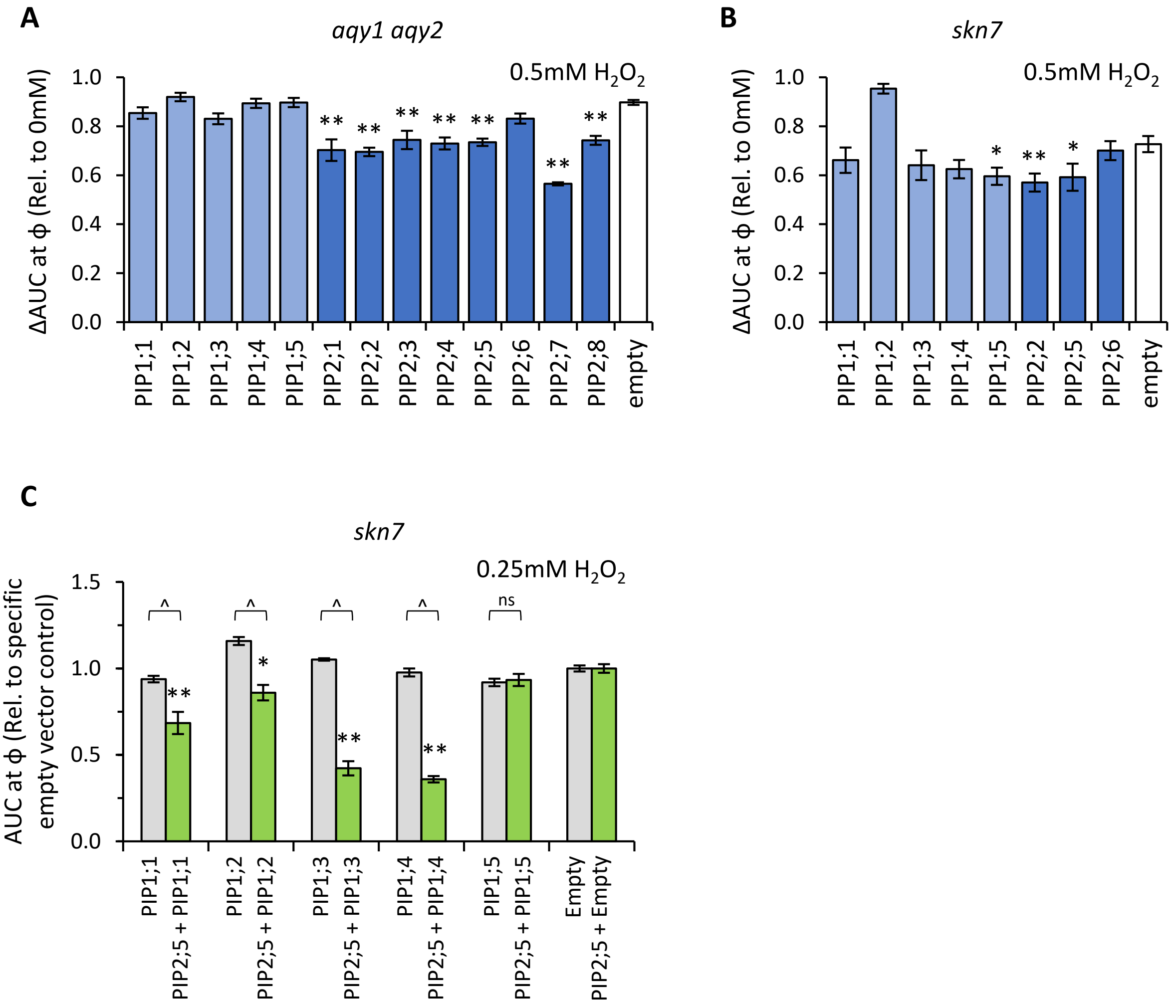


**Supplemental Figure S6. $H_2O_2$ permeability assays. A**, Relative AUC for *aqy1 aqy2* yeast expressing each AtPIP gene exposed to 0.5mM $H_2O_2$. **D**, Relative AUC for *skn7* yeast expressing *AtPIP* genes exposed to 0.5mM $H_2O_2$. **C**, Relative AUC for *skn7* yeast exposed to 0.25mM $H_2O_2$ expressing *AtPIP1* singly (grey) or together with *AtPIP2;5* (green). Each set is standardized to their respective empty vector control. All error bars are SEM. For A and B, asterisks indicate statistical difference from empty vector control, ANOVA with Fishers LSD test (* $P < 0.05$; ** $P < 0.01$). For C, asterisks indicate statistical difference from respective empty vector control, ANOVA with Fishers LSD test (* $P < 0.05$; ** $P < 0.01$); chevrons (^) indicate statistical difference between single vs. co-expression (Student's *t* test $P < 0.01$). N = 6 across 3 experimental runs for A. N = 8 across 4 experimental runs for B. For C, N = 12 across 6 experimental runs for single expressed AtPIPs and N = 6 across 3 experimental runs for co-expressed lines.

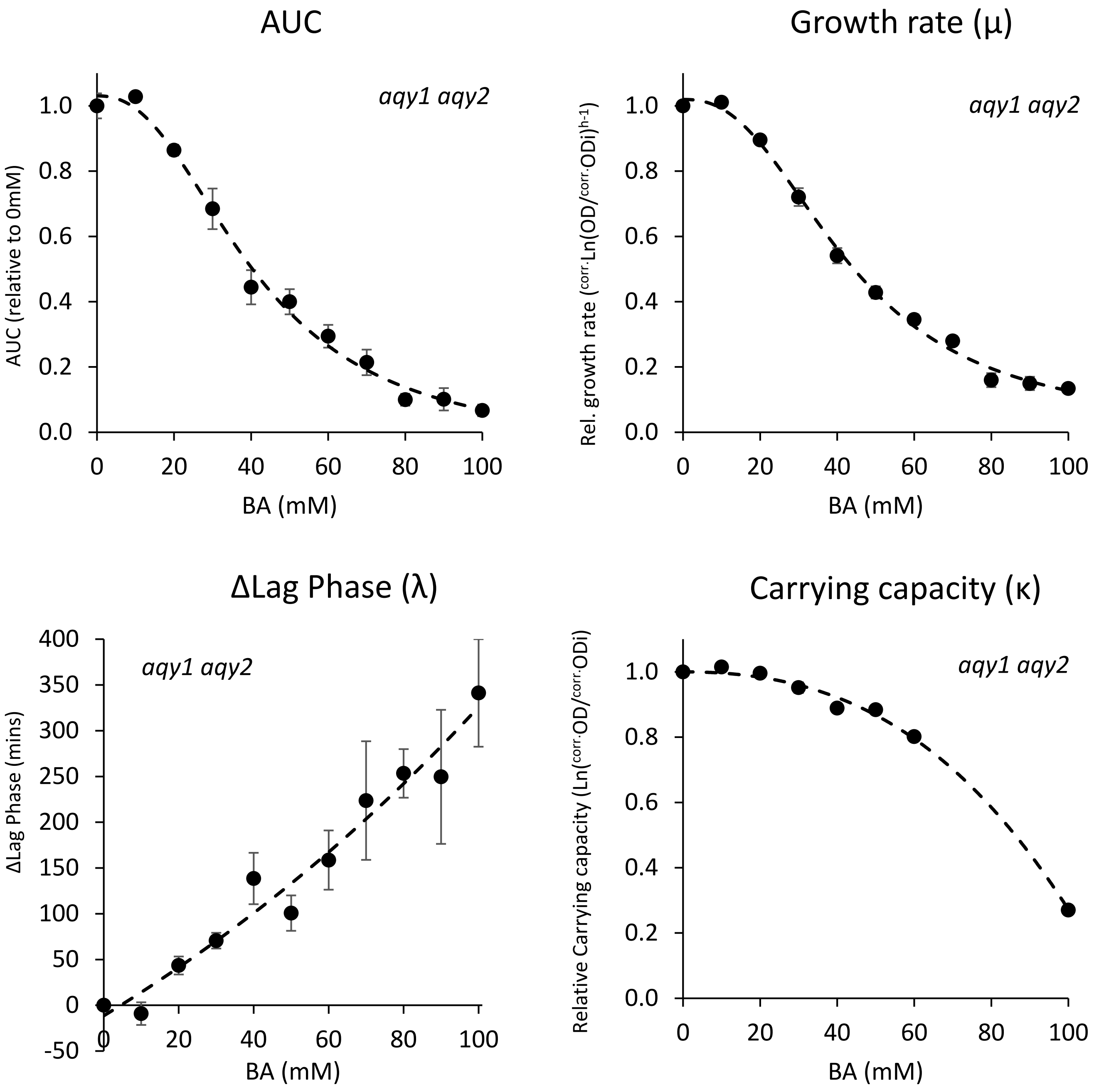


Logistic curve: $y = \dfrac{A_1 - A_2}{1 + (x / x_0)^p} + A_2$ Cubic curve: $y = A + Bx + Cx^2 + Dx^3$

**Supplemental Figure S7. Calibrating boric acid treatments for our yeast growth assay.** Growth responses of *aqy1 aqy2* yeast carrying empty vector control to increasing concentrations of boric acid (BA) added to the growth medium. BA treatments mainly reduced the rate of growth (µ). The growth responses captured by AUC was best fitted with a logistics curve. Curve fitting of individual traits: µ - logistic curve; λ – logistic curve; κ - cubic curve.

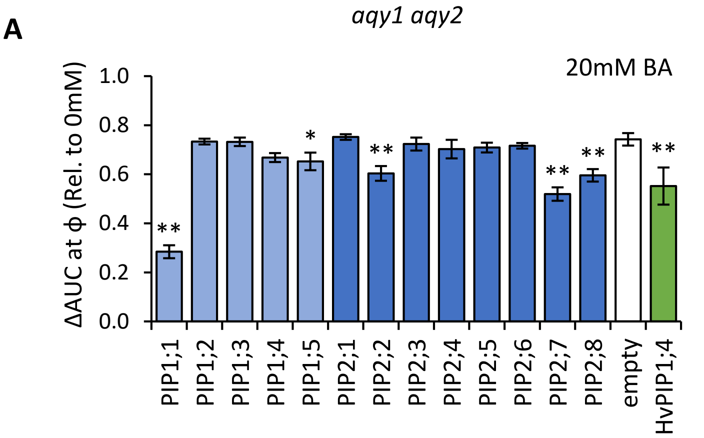


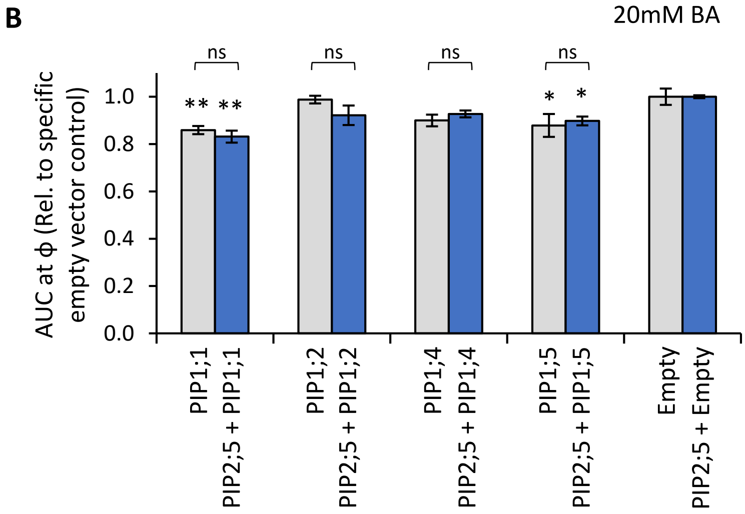


**Supplemental Figure S8. Boric acid permeability assays. A**, Relative AUC for *aqy1 aqy2* yeast expressing each *AtPIP* gene exposed to 20mM boric acid, with *HvPIP1;4* as a boric acid permeable control. **B**, Relative AUC for *aqy1 aqy2* yeast expressing *AtPIP1* singly (grey) or together with *AtPIP2;5* (blue) at 20mM boric acid. Each set is standardized to their respective empty vector control. All error bars are SEM. For A, asterisks indicate statistical difference from empty vector control, ANOVA with Fishers LSD test (* $P < 0.05$; ** $P < 0.01$). For B, asterisks indicate statistical difference from respective empty vector control, ANOVA with Fishers LSD test (* $P < 0.05$; ** $P < 0.01$. For A and B, N = 6 across 3 experimental runs.

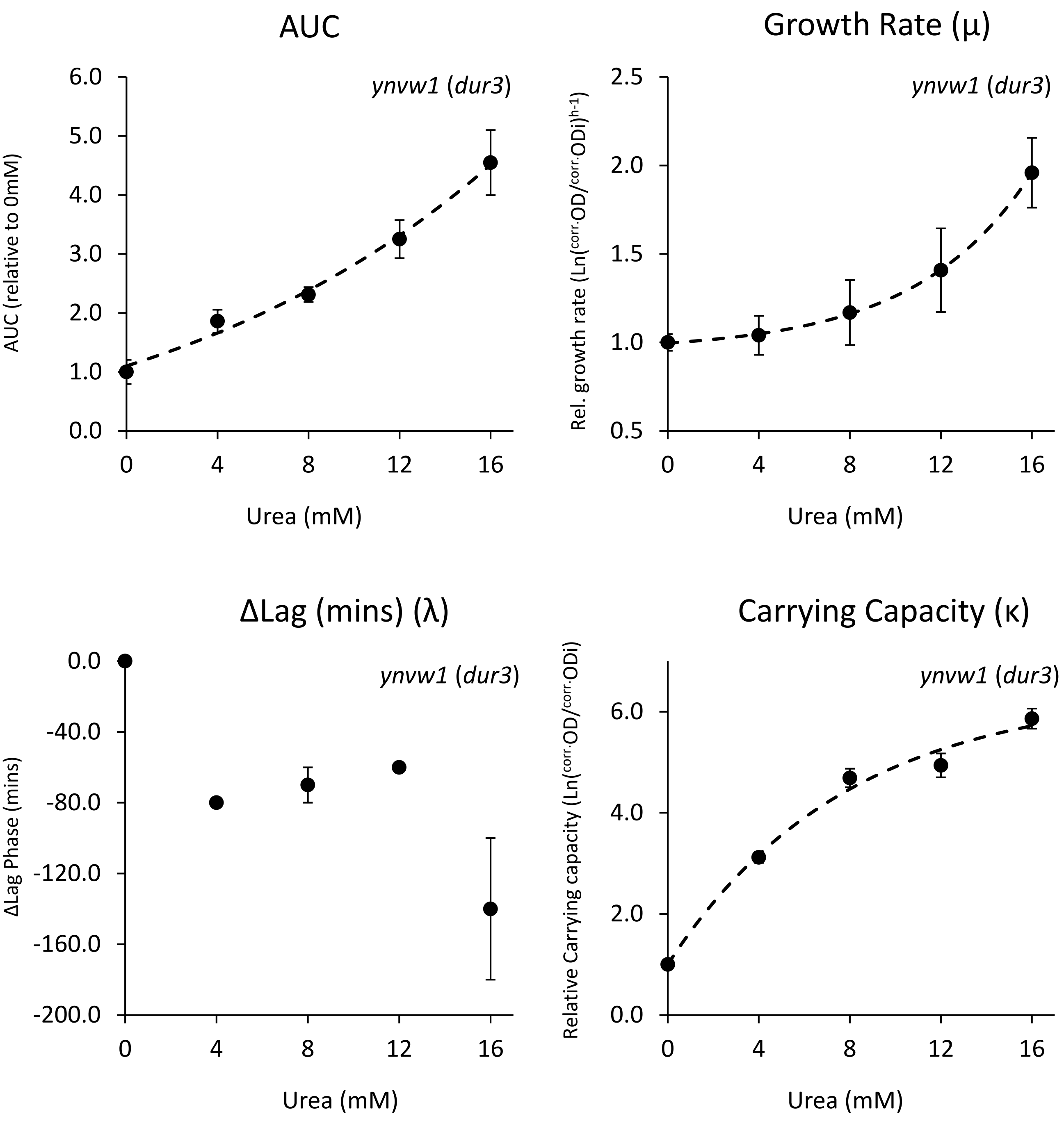


Exponential curve: $y = y_0 + Ae^{R_0 x}$

**Supplemental Figure S9. Calibrating urea treatments for yeast growth assay.** Growth responses of *ynvw1* (*dur3*) yeast carrying empty vector control to increasing concentrations of urea added to the otherwise nitrogen free growth medium. Urea supplementation mainly improves μ and κ. The growth response captured by AUC was best fitted with a exponential curve. Curve fitting of individual traits: μ - exponential curve; λ – no fit; κ - exponential curve.

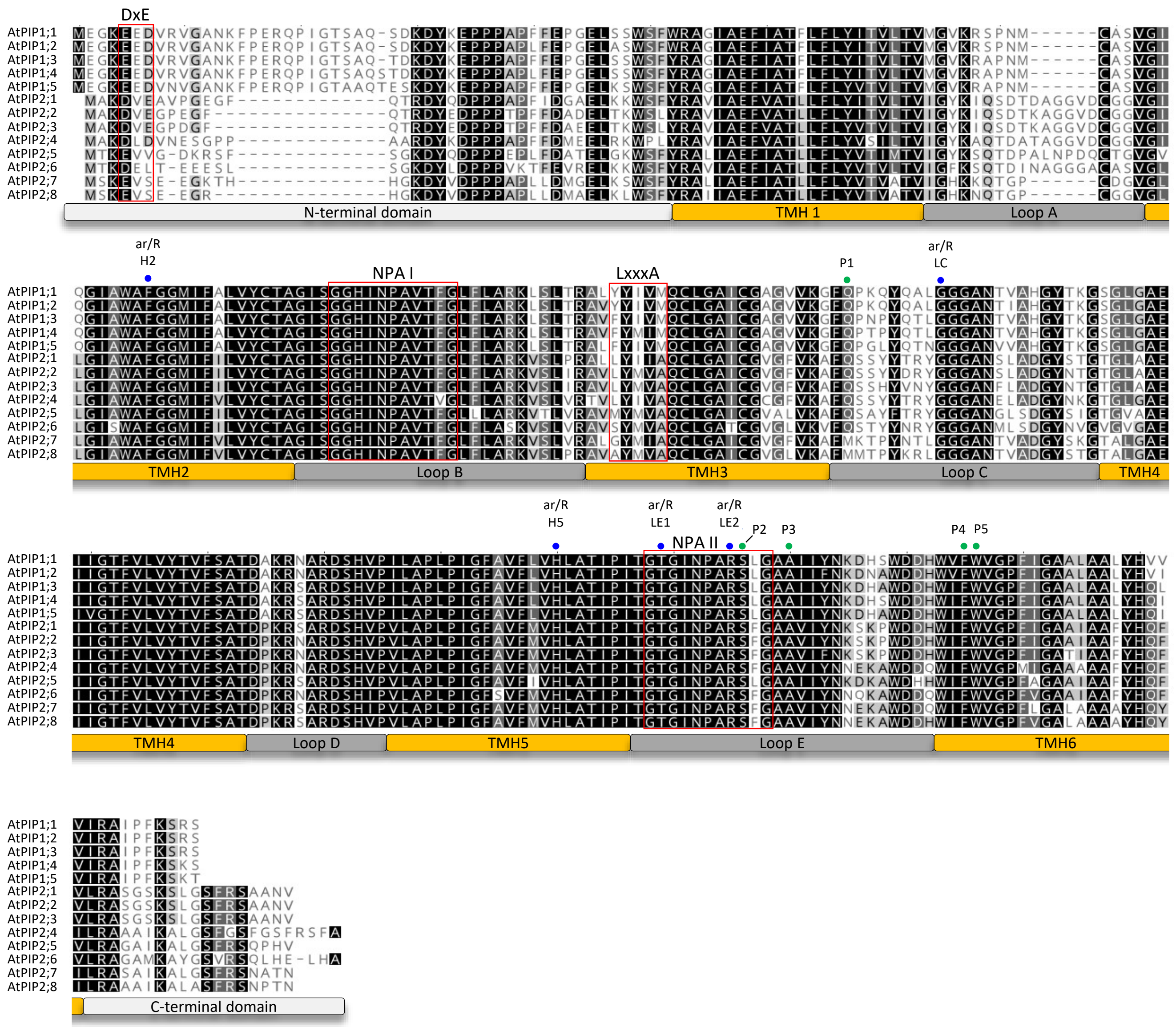

**Supplemental Figure S10. AtPIP family protein sequence alignment.** MUSCLE aligned sequences of all 13 AtPIP isoforms. The Transmembrane Helix domains (TMH1 to 6), Loop domains (Loop A-E) and N- and C-terminal domains are depicted below the sequences. Classic motifs that define substrate specificity (NPA motifs – red boxes; ar/R constriction point residues – blue dots; and Froger's positions P1-5 – green dots) and those that facilitate ER to PM trafficking (LxxA and DxE - red boxes) are labelled accordingly (also see Supplementary Table 2).

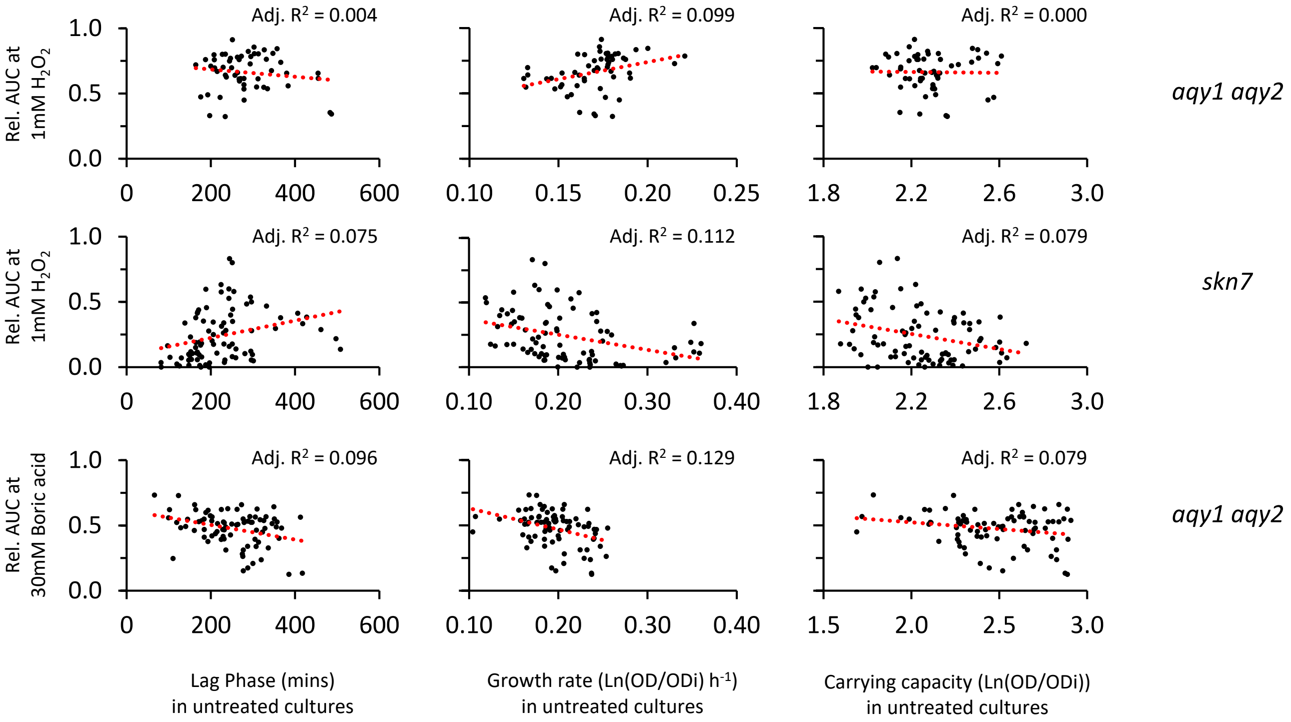


**Supplemental Figure S11. Correlation analysis examining AtPIP induced changes in inherent yeast growth characteristics and possible indirect effects on response to treatments.** No correlations were observed between treated ΔAUC values when compared against values for λ, μ, and κ from untreated cultures. This indicates that the differential growth responses of certain *AtPIP* yeast lines to specific treatments appears the direct result of AtPIP enhancement of membrane permeability to the tested substrate and not due to indirect effects through subtle changes to inherent growth characteristics.

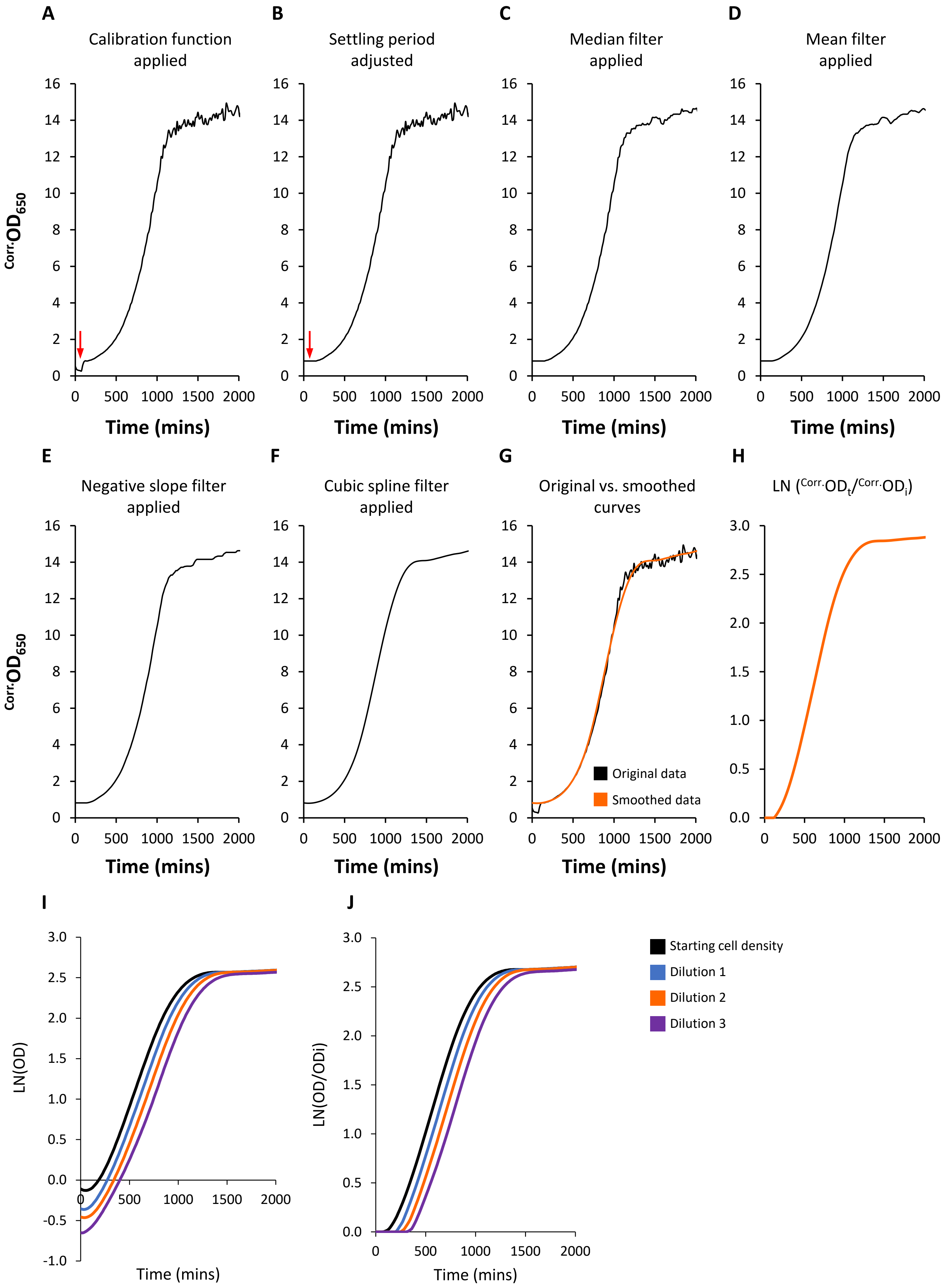


**Supplemental Figure S12. Growth curve processing. A-F**, Example showing the progressive application of filters to smooth the extracted growth curve data obtained from the micro-cultivation cultures. **G**, Overlay of the original corrected OD data and the smoothed data. **H**, The smoothed growth curve is log transformed (LN) and usually plotted as Ln($^{Corr.}OD_t$ / $^{Corr.}OD_i$). **I-J**, Toggling transformed data between LN ($^{Corr.}OD$) and LN($^{Corr.}OD_t$ / $^{Corr.}OD_i$).

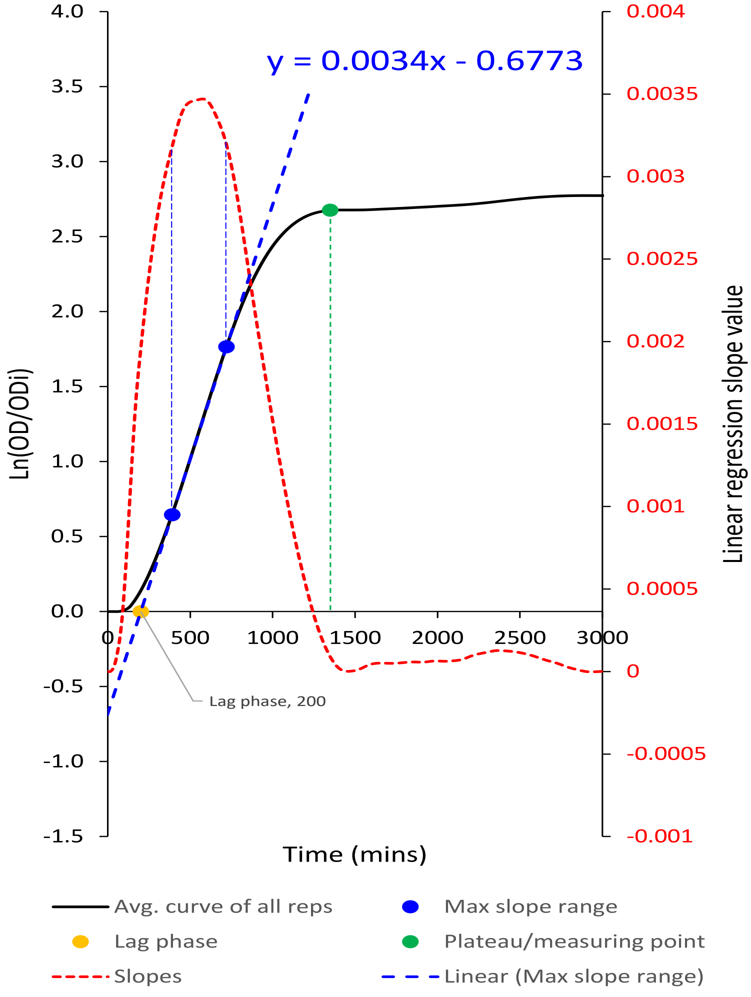


**Supplemental Figure S13. Deriving μ, λ, and κ, from a processed growth curve.**

| Gene Name | TAIR Gene ID | Protein length (aa) | G+C nucleotide content (%) | Codon Adaptation Index value | | Kozak related features | Growth characteristics of AtPIP expressing yeast | | |
|---|---|---|---|---|---|---|---|---|---|
| | | | | *in Arabidopsis thaliana* | *in Saccharomyces cerevisiae* | Nucleotide sequence surrounding mRNA AUG | Lag time (λ) (mins) | Max. growth rate (μ) $Ln(^{Corr.}OD_t / ^{Corr.}OD_i)\ h^{-1}$ | Carrying capacity (κ) Max. $Ln(^{Corr.}OD_t / ^{Corr.}OD_i)$ |
| *AtPIP1;1* | AT3G61430 | 286 | 50 | 0.80 | 0.72 | AGA ACC **AUG** GAA | 259 ± 24 $^{b,c,d}$ | 0.189 ± 0.006 $^{b,c}$ | 2.40 ± 0.05 $^{a,b,c}$ |
| *AtPIP1;2* | AT2G45960 | 286 | 50 | 0.80 | 0.71 | AGA ACC **AUG** GAA | 251 ± 14 $^{b,c,d}$ | 0.184 ± 0.002 $^{b,c}$ | 2.40 ± 0.06 $^{a,b,c}$ |
| *AtPIP1;3* | AT1G01620 | 286 | 52 | 0.76 | 0.69 | AGA ACC **AUG** GAA | 262 ± 15 $^{a,b,c,d}$ | 0.195 ± 0.010 $^{a,b,c}$ | 2.29 ± 0.05 $^{c,d}$ |
| *AtPIP1;4* | AT4G00430 | 287 | 51 | 0.78 | 0.70 | AGA ACC **AUG** GAA | 250 ± 16 $^{b,c,d}$ | 0.177 ± 0.003 $^{c,d}$ | 2.22 ± 0.03 $^{d}$ |
| *AtPIP1;5* | AT4G23400 | 287 | 49 | 0.79 | 0.72 | AGA ACC **AUG** GAA | 250 ± 19 $^{b,c,d}$ | 0.178 ± 0.002 $^{c,d}$ | 2.36 ± 0.05 $^{b,c,d}$ |
| *AtPIP2;1* | AT3G53420 | 287 | 50 | 0.76 | 0.71 | AGA ACC **AUG** GCA | 265 ± 12 $^{a,b,c,d}$ | 0.183 ± 0.005 $^{b,c}$ | 2.47 ± 0.06 $^{a,b}$ |
| *AtPIP2;2* | AT2G37170 | 285 | 51 | 0.73 | 0.68 | AGA ACC **AUG** GCC | 240 ± 13 $^{c,d}$ | 0.199 ± 0.012 $^{a,b}$ | 2.53 ± 0.08 $^{a}$ |
| *AtPIP2;3* | AT2G37180 | 285 | 51 | 0.73 | 0.67 | AGA ACC **AUG** GCT | 225 ± 14 $^{d}$ | 0.181 ± 0.007 $^{b,c,d}$ | 2.49 ± 0.07 $^{a,b}$ |
| *AtPIP2;4* | AT5G60660 | 291 | 51 | 0.73 | 0.69 | AGA ACC **AUG** GCA | 257 ± 24 $^{b,c,d}$ | 0.164 ± 0.005 $^{d}$ | 2.36 ± 0.07 $^{a,b,c,d}$ |
| *AtPIP2;5* | AT3G54820 | 286 | 52 | 0.76 | 0.70 | AGA ACC **AUG** ACG | 228 ± 31 $^{c,d}$ | 0.140 ± 0.008 $^{e}$ | 2.01 ± 0.07 $^{e}$ |
| *AtPIP2;6* | AT2G39010 | 289 | 51 | 0.74 | 0.68 | AGA ACC **AUG** ACG | 308 ± 34 $^{a,b}$ | 0.187 ± 0.005 $^{b,c}$ | 2.53 ± 0.07 $^{a}$ |
| *AtPIP2;7* | AT4G35100 | 280 | 50 | 0.81 | 0.74 | AGA ACC **AUG** TCG | 315 ± 24 $^{a}$ | 0.212 ± 0.008 $^{a}$ | 2.49 ± 0.05 $^{a,b}$ |
| *AtPIP2;8* | AT2G16850 | 278 | 49 | 0.78 | 0.73 | AGA ACC **AUG** TCA | 291 ± 25 $^{a,b,c}$ | 0.201 ± 0.012 $^{a,b}$ | 2.41 ± 0.07 $^{a,b,c}$ |
| empty vector | - | - | | - | - | - | 279 ± 20 $^{a,b,c,d}$ | 0.195 ± 0.006 $^{a,b}$ | 2.51 ± 0.05 $^{a,b}$ |

Low Mid High

**Supplemental Table S1. *AtPIP* codon compatibility for heterologous expression in yeast and growth characteristics of *AtPIP* expressing yeast lines.** The AtPIPs have similar good codon compatibility (CAI values and GC contents) for heterologous expression in yeast, with yeast CAI values similar to native yeast *AQP* genes (*AQY1*: 0.71 CAI and 51% G+C; *AQY2*: 0.79 CAI and 46% G+C). The upstream Kozak region (red nucleotides) were engineered as part of the vector construction. AtPIP expressing yeast cultured in standard selection medium (no treatment) show slight differences in growth characteristics between each other and empty vector control. Superscript letters denote statistical groupings, ANOVA with Tukey's test ($P < 0.05$). N ≥ 10 for each line.

| | | | Length of protein domain (aa) | | | | | | | | | | | | | NPA I motif | NPA II motif | ar/R | | | | | Froger's positions | | | | | PM targeting | | C-terminal domain |
|---|---|---|---|---|---|---|---|---|---|---|---|---|---|---|---|---|---|---|---|---|---|---|---|---|---|---|---|---|---|---|
| Protein Name | TAIR ID | Protein length (aa) | N-term | TM1 | LA | TM2 | LB | TM3 | LC | TM4 | LD | TM5 | LE | TM6 | C-term | LB | LE | H2 | LC | H5 | LE1 | LE2 | P1 | P2 | P3 | P4 | P5 | DxE | LxxxA | |
| AtPIP1;1 | AT3G61430 | 286 | 51 | 21 | 13 | 21 | 25 | 21 | 23 | 21 | 12 | 21 | 26 | 21 | 10 | GGHI**NPA**VTFG | GTGI**NPA**RSLG | F | G | H | T | R | Q | S | A | F | W | EED | **Y**YIV**M** | IRAIPFK**S**R**S** |
| AtPIP1;2 | AT2G45960 | 286 | 51 | 21 | 13 | 21 | 25 | 21 | 23 | 21 | 12 | 21 | 26 | 21 | 10 | GGHI**NPA**VTFG | GTGI**NPA**RSLG | F | G | H | T | R | Q | S | A | F | W | EED | **Y**YIV**M** | IRAIPFK**S**R**S** |
| AtPIP1;3 | AT1G01620 | 286 | 51 | 21 | 13 | 21 | 25 | 21 | 23 | 21 | 12 | 21 | 26 | 21 | 10 | GGHI**NPA**VTFG | GTGI**NPA**RSLG | F | G | H | T | R | Q | S | A | F | W | EED | **F**YIV**M** | IRAIPFK**S**R**S** |
| AtPIP1;4 | AT4G00430 | 287 | 52 | 21 | 13 | 21 | 25 | 21 | 23 | 21 | 12 | 21 | 26 | 21 | 10 | GGHI**NPA**VTFG | GTGI**NPA**RSLG | F | G | H | T | R | Q | S | A | F | W | EED | **F**YMI**M** | IRAIPFK**S**K**S** |
| AtPIP1;5 | AT4G23400 | 287 | 52 | 21 | 13 | 21 | 25 | 21 | 23 | 21 | 12 | 21 | 26 | 21 | 10 | GGHI**NPA**VTFG | GTGI**NPA**RSLG | F | G | H | T | R | Q | S | A | F | W | EED | **F**YIV**M** | IRAIPFK**S**KT |
| AtPIP2;1 | AT3G53420 | 287 | 38 | 21 | 20 | 21 | 25 | 21 | 23 | 21 | 12 | 21 | 26 | 21 | 18 | GGHI**NPA**VTFG | GTGI**NPA**RSFG | F | G | H | T | R | Q | S | A | F | W | DVE | **L**YII**A** | LRA**S**G**S**K**S**LG**S**FR**S**AANV |
| AtPIP2;2 | AT2G37170 | 285 | 36 | 21 | 20 | 21 | 25 | 21 | 23 | 21 | 12 | 21 | 26 | 21 | 18 | GGHI**NPA**VTFG | GTGI**NPA**RSFG | F | G | H | T | R | Q | S | A | F | W | DVE | **L**YMV**A** | LRA**S**G**S**K**S**LG**S**FR**S**AANV |
| AtPIP2;3 | AT2G37180 | 285 | 36 | 21 | 20 | 21 | 25 | 21 | 23 | 21 | 12 | 21 | 26 | 21 | 18 | GGHI**NPA**VTFG | GTGI**NPA**RSFG | F | G | H | T | R | Q | S | A | F | W | DVE | **L**YMV**A** | LRA**S**G**S**K**S**LG**S**FR**S**AANV |
| AtPIP2;4 | AT5G60660 | 291 | 38 | 21 | 20 | 21 | 25 | 21 | 23 | 21 | 12 | 21 | 26 | 21 | 22 | GGHI**NPA**VTVG | GTGI**NPA**RSFG | F | G | H | T | R | Q | S | A | F | W | DLD | **L**YIV**A** | RAAAIKALG**S**FG**S**FG**S**FR**S**FA |
| AtPIP2;5 | AT3G54820 | 286 | 37 | 21 | 20 | 21 | 25 | 21 | 23 | 21 | 12 | 21 | 26 | 21 | 18 | GGHI**NPA**VTFG | GTGI**NPA**RSLG | F | G | H | T | R | Q | S | A | F | W | EEV | **M**YMV**A** | RAGAIKALG**S**FR**S**QPHV |
| AtPIP2;6 | AT2G39010 | 289 | 37 | 21 | 20 | 21 | 25 | 21 | 23 | 21 | 12 | 21 | 26 | 21 | 21 | GGHI**NPA**VTFG | GTGI**NPA**RSFG | F | G | H | T | R | Q | S | A | F | W | DEL | **S**YMV**A** | RAGAMKAYG**S**VR**S**QLHELHA |
| AtPIP2;7 | AT4G35100 | 280 | 37 | 21 | 14 | 21 | 25 | 21 | 23 | 21 | 12 | 21 | 26 | 21 | 18 | GGHI**NPA**VTFG | GTGI**NPA**RSFG | F | G | H | T | R | M | S | A | F | W | EVS | **G**YMI**A** | RA**S**AIKALG**S**FR**S**NATN |
| AtPIP2;8 | AT2G16850 | 278 | 35 | 21 | 14 | 21 | 25 | 21 | 23 | 21 | 12 | 21 | 26 | 21 | 18 | GGHI**NPA**VTFG | GTGI**NPA**RSFG | F | G | H | T | R | M | S | A | F | W | EVS | **A**YMV**A** | RAAAIKALA**S**FR**S**NPTN |

**Supplemental Table S2. Protein domain lengths and amino acid composition of AtPIPs at known substrate selectivity positions and other important motifs.** The N-terminal domain of AtPIP1s is distinctly longer than those of AtPIP2s, whereas the C-terminal domain is longer in AtPIP2s than AtPIP1s. More serine residues are present in the C-terminal domain of AtPIP2s than AtPIP1s (bold black), with two positions in the AtPIP2s likely to be phosphorylation targets (bold blue). The extracellular loop A domain is longer in AtPIP2;1 to 2;6 isoforms. Classic motifs that define substrate specificity (i.e. NPA motifs, ar/R constriction point, and Froger's positions) are nearly identical, with only seemingly minor conserved differences. There is substantial variation in the motifs associated with PM targeting between the AtPIP isoforms.
